# Grazing versus mowing: productivity and forage quality of mixtures and cultivars of perennial ryegrass

**DOI:** 10.64898/2026.10.15.759330

**Authors:** Damian Käch, Daniel Suter, Bruno Studer, Roland Kölliker, Beat Reidy

## Abstract

Perennial ryegrass (*Lolium perenne* L.) is an important grass species in temporary grasslands, where it is often sown as a mixture of multiple cultivars. In variety testing, however, candidate cultivars are evaluated as single-cultivar stands under standardised mechanical defoliation. In practice, swards are mown, grazed, or both, where grazing adds treading, selective defoliation, and heterogeneous nutrient return. Whether cultivar rankings obtained under mechanical defoliation also hold under grazing, and whether mixtures perform differently than their component cultivars, are unresolved. We compared six early-heading perennial ryegrass cultivars (three diploid, three tetraploid), and three 50:50 two-cultivar mixtures. The plots were grown under three defoliation treatments (i.e., regular cutting, simulated grazing, and grazing by dairy cows), over three years at two sites. Here we show that defoliation treatment and year affected dry matter yield (DMY) and forage quality stronger than the cultivar grown. The effect of ploidy depended on the defoliation treatment. Tetraploid cultivars had slightly higher DMY under regular cutting and simulated grazing and higher metabolisable energy throughout, whereas diploid cultivars had slightly higher DMY under grazing. Simulated grazing matched the forage quality of grazed swards, including cultivar ranking, but not the cultivar ranking for DMY. Forage quality rankings can therefore be assessed through simulated grazing, whereas DMY rankings of cultivars intended for grazing require evaluation under grazing. Diploid-tetraploid mixtures were intermediate in forage quality and, under grazing, slightly higher in DMY when compared to their individual components. Evaluating cultivars and mixtures under grazing would better reflect their relative performance in the field.

## 1 Introduction

Grasslands are the basis for sustainable dairy and beef production. They cover 43% of the agricultural area in Western Europe, and even more than 70% in some countries such as Switzerland and Ireland (Boch et al., 2020). Much of this area is not suitable for arable cropping, meaning that grassland-based ruminant production converts human-inedible biomass into milk and meat without competing for land that could feed people directly (Leiber, 2022; Mottet et al., 2017; van Zanten et al., 2018). Beyond producing forage, grasslands supply a range of ecosystem services, including biodiversity conservation and carbon storage (Bengtsson et al., 2019).

Perennial ryegrass (*Lolium perenne* L.) is an important grass species in temporary grasslands of temperate regions (Wilkins and Humphreys, 2003). Breeding targets such as a high dry matter yield (DMY), a high forage quality, and persistence under repeated defoliation have long been pursued (Faville et al., 2020). Candidate cultivars are evaluated in variety testing, a multi-year, multi-site assessment of the above and additional traits under standardised defoliation treatments, which forms the basis for the national lists of recommended varieties (e.g., Suter et al., 2025). Perennial ryegrass is a natural diploid, but is also bred as tetraploid cultivars after artificial chromosome duplication. Tetraploid cultivars have larger cells and higher cell content-to-cell wall ratio and therefore a higher forage quality, whereas diploid cultivars form denser swards with more and smaller tillers (Beecher et al., 2015; Griffiths et al., 2016; Sugiyama, 2005). These contrasting morphologies suggest that diploid and tetraploid cultivars are not equally suited to all defoliation treatments, but this interaction between ploidy and defoliation treatment is rarely quantified within a single experiment.

Forage quality is not a single trait but reflects the balance between readily available cell content (e.g., crude protein and non-fibre carbohydrates) and cell wall components (i.e., structural carbohydrates and lignin). These components together determine digestibility and metabolisable energy, and thus intake and animal performance (Bruinenberg et al., 2002; Terry and Tilley, 1964). Although cultivar differences in forage quality are heritable and targeted in breeding (Arojju et al., 2020), the balance between cell content and cell wall components shifts strongly with plant maturity and is therefore highly sensitive to how frequently and how low a sward is defoliated. Cultivar differences in forage quality can consequently only be interpreted in the context of the defoliation treatment under which they were assessed.

In practice, grasslands are mown, grazed, or used in a combination of both. The two defoliation treatments differ in defoliation height and frequency, and in the pressure they exert on the sward: mowing removes the herbage uniformly in height, timing, and across the plot area, whereas grazing adds treading, selective defoliation, and heterogeneous nutrient return through excreta. Grazing therefore represents a distinct selection pressure, and tolerance to it is a breeding target on its own. During cultivar development, however, swards are evaluated almost exclusively under mechanical defoliation. To approximate grazing without animals, official variety testing in countries such as the United Kingdom and Switzerland applies simulated grazing, a frequent mechanical cutting at a low defoliation height (i.e., up to 10 defoliations per year at *≈* 4 cm), alongside regular cutting, a less frequent cutting at a greater height (i.e., *≈* 5 defoliations per year at *≈* 8 cm). In Switzerland, simulated grazing is not intended to estimate the yield of grazed swards but to assess the tillering of cultivars under frequent, low defoliation. Whether it also reproduces the cultivar ranking for DMY and forage quality of grazed swards is unresolved: most studies compared values, not rankings, and reported similar to higher DMY and similar forage quality under simulated grazing, with high variability between studies attributed to the inclusion of animals (Binnie and Chestnutt, 1991; Cashman et al., 2016; Grace et al., 2019; Jackson and Williams, 1979). Grazing experiments are therefore essential to quantify the effect of animals and to close the knowledge gap between cultivar development and on-farm practice.

Another knowledge gap is that candidate cultivars are evaluated as single-cultivar stands, although they are later sown in mixtures (Wilkins and Humphreys, 2003). Mixing cultivars can raise and stabilise yield and improve tolerance of plant antagonists, in a similar way that mixing species does at the between-species level (Haughey et al., 2018; Küchenmeister et al., 2012; Meilhac et al., 2019; Nyfeler et al., 2009; Pollnac et al., 2014; Prieto et al., 2015; Wan et al., 2022). If diploid and tetraploid cultivars are suited to different defoliation treatments, mixing them could result in seed mixtures that perform adequately under both mowing and grazing. However, evidence on how cultivar mixtures perform relative to their components is scarce and ambivalent, ranging from a higher DMY of the mixtures to a performance intermediate between the corresponding cultivars (Gilliland et al., 2011; Jones and Roberts, 1994; Rhodes, 1970; Trenbath, 1974; Tubritt et al., 2021). Moreover, the composition of such mixtures may shift over time, as the cultivar better adapted to a given defoliation treatment gains in abundance (Gilliland et al., 2011). Such shifts matter for breeders, because they reflect persistence and competitiveness. They can be inferred indirectly by comparing the performance of the mixture to that of the corresponding cultivars grown as pure stands or quantified directly by identifying individual cultivars within the mixed sward using allozyme or molecular markers (Kennedy et al., 1985; van der Laat et al., 2025; Liu et al., 2010). However, studies applying these methods to compare compositional shifts in cultivar mixtures across contrasting cutting and grazing defoliation treatments remain rare.

We therefore assessed DMY and forage quality of diploid and tetraploid perennial ryegrass cultivars and of mixtures of two cultivars. The plots were managed under regular cutting, simulated grazing, and grazing by dairy cows, over three years at two sites. We hypothesised that (i) DMY and forage quality are more strongly affected by defoliation treatment and year than by the cultivar; (ii) diploid and tetraploid cultivars perform differently depending on the defoliation treatment; (iii) mixtures perform intermediate relative to their component cultivars and shift in composition over time; and (iv) simulated grazing is a suitable proxy for the cultivar ranking in DMY and forage quality under grazing.

## 2 Materials and methods

### 2.1 Field experiment

The field experiment was established in the autumn of 2019 on two farms in the central plateau of Switzerland, Langenthal BE [47*^◦^*13*^′^*3.40*^′′^*N, 7*^◦^*48*^′^*15.24*^′′^*E] and Eschenbach LU [47*^◦^*8*^′^*16.54*^′′^*N, 8*^◦^*18*^′^*16.89*^′′^*E]. Over the study period (2019-2023), mean annual air temperature was 10.3°C at Langenthal and 10.7°C at Eschenbach, with monthly means ranging from 0.2 to 20.6°C and 0.4 to 21.3°C, respectively (Supplementary Figure S1, Supplementary Table S3). Mean annual precipitation was 1042.6 mm at Langenthal and 1206.5 mm at Eschenbach. Within years, the two locations differed in the distribution of precipitation: Precipitation in Langenthal was higher in summer and lower in winter when compared to Eschenbach.

Nine plot compositions based on six early heading *Lolium perenne* L. cultivars were sown in drills at a sowing density of 500 germinable seeds m^-2^ (Table 1). Three cultivars were diploid (‘Arara‘ [Ara], ‘Araias‘ [Ari], ‘Repentinia‘ [Rep]) and three tetraploid (‘Artonis‘ [Art], ‘Arcturus‘ [Arc], ‘Algira‘ [Alg]; Käch et al., 2026). Six plot compositions were composed of a single cultivar, whereas three contained two cultivars sown at a 50:50-ratio of germinable seeds, where each mixture contained Ari and either Rep (Rep50), Art (Art50) or Alg (Alg50).

**Table 1:**
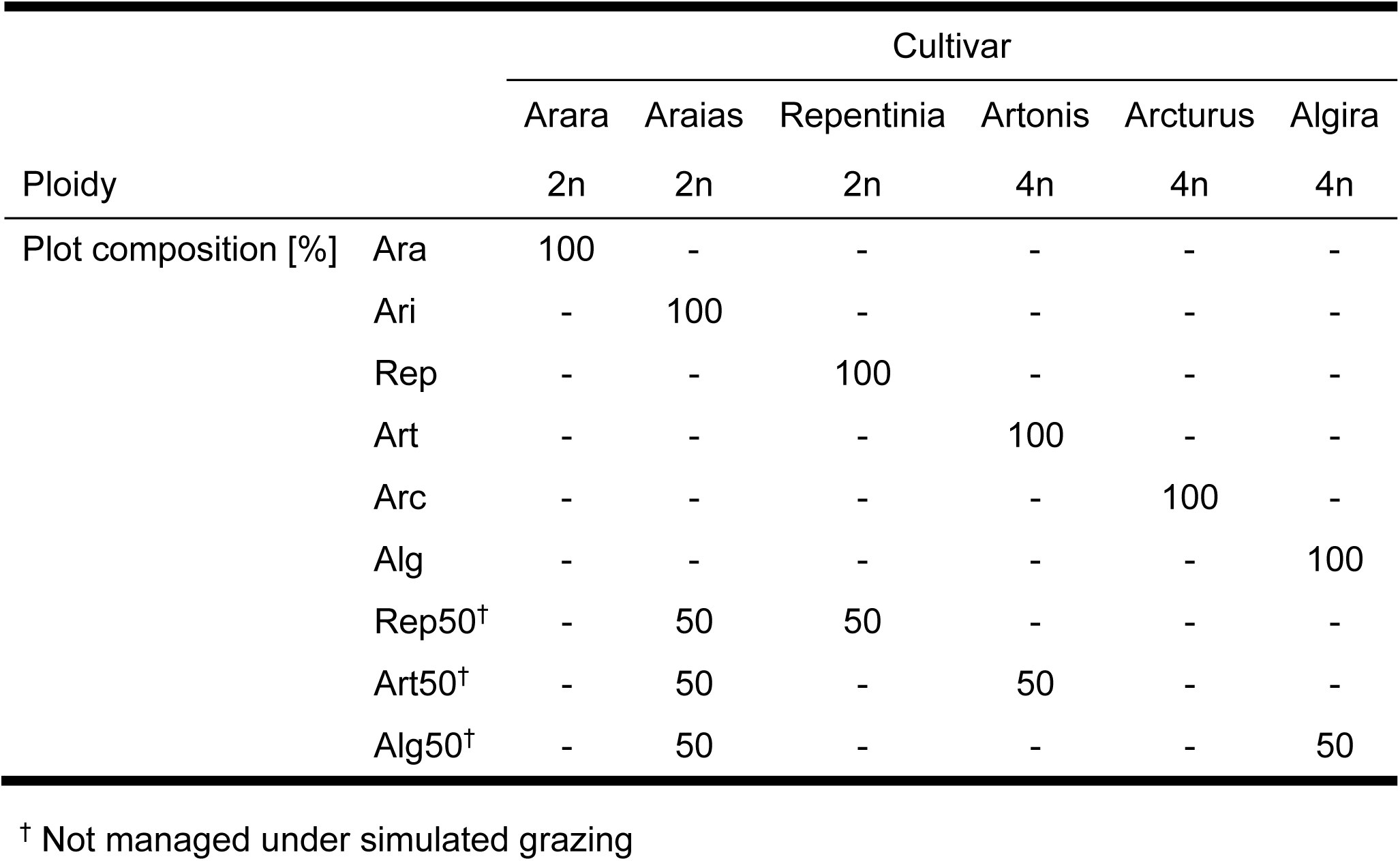
Plot compositions based on six early-heading cultivars of *Lolium perenne* L.

Three defoliation treatments were applied: regular cutting with five defoliations per year at a mowing height of 8 cm, grazing by 30-40 cows ha^-1^ during two consecutive half-days per defoliation with seven to nine defoliations per year (depending on the practice of the farmers) to a target post-grazing height of 4 cm, and simulated grazing with eight defoliations per year at a mowing height of 4 cm. The high stocking density under grazing aimed to reduce selective grazing. While the swards under regular cutting and simulated grazing were sown in plots of 1.5 by 6 m, those under grazing were sown in plots of 6 by 6 m. Plots under regular cutting and simulated grazing were fenced in, whereas those under grazing were part of a larger pasture. All plots were arranged in a split-plot design in three replicates, with the mixtures (Rep50, Art50, and Alg50) not being grown under simulated grazing.

Plots under regular cutting and simulated grazing were fertilised according to the practice in breeding and variety testing (i.e., 260 kg N ha^-1^ a^-1^) using ammonium nitrate after each defoliation event. The plots under grazing were fertilised according to the practice of the farmers (i.e., 160 kg N ha^-1^ a^-1^, not accounting for excretion during grazing) using manure at the beginning of the vegetation period and ammonium nitrate and phosphate during the vegetation period. In addition to these fertiliser applications, the excretion during grazing is estimated to account for 80 kg N ha^-1^ a^-1^ (Richner and Sinaj, 2017; Supplementary Section A). In total, this corresponded to a total fertiliser application of approximately 240 kg N ha^-1^ a^-1^ for the plots under grazing. The manure used for fertilising was sampled in spring and autumn to measure the nutrient content. Additionally, evenly distributed soil samples were taken before establishing the experiment and in the spring of 2022 to measure soil characteristics and content (Supplementary Figure S2). Manure and soil samples were analysed by Eric Schweizer AG (Thun, Switzerland; Supplementary Section A, Supplementary Table S4, Supplementary Table S5). Additionally, weather data were taken from the stations Eschenbach (Canton Lucerne network) and Langenthal (DTN Schweiz AG, Appenzell, Switzerland) and downloaded via the portal IDAWEB of MeteoSwiss (Kloten, Switzerland).

### 2.2 Dry matter yield

After the year of establishment, DMY was measured in 2021, 2022, and 2023, i.e., in the second to fourth growing season after sowing (Supplementary Table S2). For plots under regular cutting and simulated grazing, a forage harvester (Wintersteiger, Ried im Innkreis, Austria) was used to harvest and weigh the fresh matter. Plots under simulated grazing were additionally mowed using a lawn mower to reach the aimed at post-mowing height. For the grazing plots, a bar mower was used to mow a strip of 0.7 by 6 m to a height of 4 cm before the grazing event, where the strip location was shifted for each grazing event. The fresh matter of the strip was weighed using a net and a hanging scale. For all plots, a representative sample was taken and dried at 105 °C for at least 24 hours to determine dry matter content and calculate DMY.

### 2.3 Forage quality

Forage quality was assessed in samples taken at the first, second, summer, and last defoliation (Supplementary Table S2). After sampling, the plant material was dried at 60 °C for at least 24 hours and milled to 1 mm. Reference values of neutral detergent fibre (NDF), acid detergent fibre (ADF) and lignin (ADL), crude protein (CP), and non-fibre carbohydrates (NFC) for 168 representative samples were determined following standard AOAC procedures (NDF: Van Soest et al., 1991; ADF: AOAC 973.18; ADL: AOAC 973.18; CP: AOAC 990.03; Dairy One, Ithaca, NY, USA; Supplementary Section A). NFC reference values were obtained by Dairy One according to their standard procedures. These reference values were used to calibrate near-infrared (NIR) spectra for prediction of the same parameters in all remaining samples (Supplementary Table S6). Metabolisable energy (ME) was calculated from these parameters, resulting in a total of six forage quality parameters (Agroscope, 2021; Supplementary Section A).

### 2.4 Data analysis

#### 2.4.1 Dry matter yield

The analysis of DMY was based on the yearly sum. The dataset contained missing values in the following plots: first defoliation of grazing plots and first two defoliations of regular cutting and simulated grazing plots in Langenthal in 2021. Since these missing values would have affected the yearly sum of DMY, they were imputed using plots with similar characteristics. Imputation was conducted on each defoliation treatment separately. The following imputation methods were tested based on predicting known values of plots that were similar to the plots to impute and comparing these to the actual values: linear regression, random forest using randomForest (Liaw and Wiener, 2002), support vector machine using e1071 (Meyer et al., 2024), and arithmetic mean of similar plots. Linear regression had the best balance between complexity and root mean square error and was used to impute missing values for all defoliation treatments.

Statistical analysis was conducted based on the following equation:

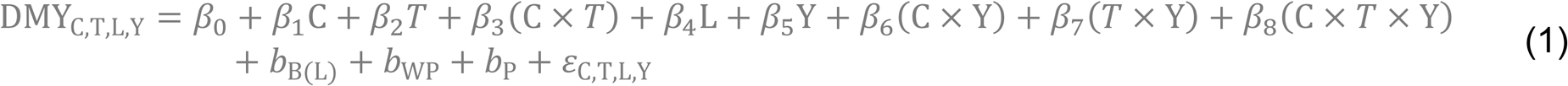

where DMY_C,T,L,Y_ is the yearly sum of DMY for each combination of plot composition (C), defoliation treatment (T), location (L), and year (Y); β_1_ to β_8_ are the coefficients of the fixed effects; b_B(L)_, b_WP_, and b_P_ are the random effects of block within location, whole plot (i.e., the defoliation-treatment-level unit to which T was randomised within each block), and plot (i.e., the subplot to which C was randomised within each whole plot), respectively; and *ε*_C,T,L,Y_ is the residual error term. Based on the fitted model using lmerTest (Kuznetsova et al., 2017) and pbkrtest (Halekoh and Højsgaard, 2014), an analysis of variance (ANOVA) was conducted, and estimated marginal means with 95% confidence intervals were computed using emmeans (Lenth, 2025).

#### 2.4.2 Forage quality

Forage quality was statistically analysed based on the following equation:

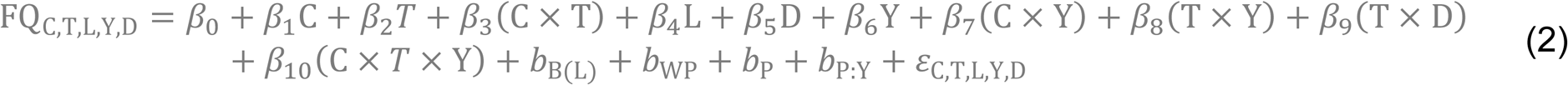

where FQ_C,T,L,Y,D_ is the forage quality parameter (CP, NFC, NDF, ADF, ADL or ME) for each combination of plot composition (C), defoliation treatment (T), location (L), year (Y), and defoliation (D), i.e. the sampled defoliation event within the year (first, second, summer, or last; Supplementary Table S2); β_1_ to β_10_ are the coefficients of the fixed effects; b_B(L)_, b_WP_, b_P_, and b_P:Y_ are the random effects of block within location, whole plot (i.e., the defoliation-treatment-level unit to which T was randomised within each block), plot (i.e., the subplot to which C was randomised within each whole plot), and plot-year (to account for the four repeated defoliation measurements within a plot-year), respectively; and *ε*_C,T,L,Y,D_ is the residual error term. In addition to the factors included in Equation 1, Equation 2 incorporates defoliation number and the random effect of plot-year. Similarly to DMY, and for each forage quality parameter, the fitted model was used for an ANOVA, and estimated marginal means with 95% confidence intervals were computed.

#### 2.4.3 Temporal shift in plot composition

The temporal shift in plot composition was estimated by comparing the mixtures and the corresponding cultivars per year using the following formula:

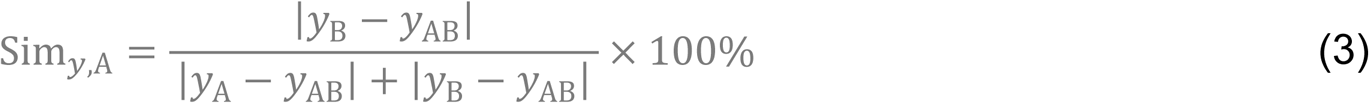

where Sim*_y,_*_A_ is the relative similarity in *y* (i.e., DMY, CP, NFC, NDF, ADF, ADL or ME) of mixture AB (i.e., Rep50, Art50 or Alg50) to cultivar A (i.e., Ari). Mixture AB contains cultivar A and cultivar B (i.e., Rep, Art or Alg). Possible values range from 0% to 100%, corresponding to complete identity with cultivar B or Ari, respectively.

#### 2.4.4 Cultivar ranking

To compare how cultivars ranked between defoliation treatments, Spearman rank correlations were computed between each pair of defoliation treatments, based on the estimated marginal means of the six cultivars per parameter (i.e., DMY, CP, NFC, NDF, ADF, ADL, and ME).

Data processing, analysis and visualisation were conducted with R v4.5.0 (R core team, 2025). A complete list of packages and versions are provided as Supporting Information (Supplementary Section A).

## 3 Results

### 3.1 Dry matter yield

The yearly sum of DMY was mainly affected by the year and defoliation treatment (*P* < 0.0001), whereas plot composition had no significant main effect (*P* > 0.05), although it interacted significantly with defoliation treatment (*P* < 0.05; Table 2, Supplementary Table S8). Regular cutting resulted in the highest DMY, closely followed by simulated grazing, whereas grazing resulted in clearly the lowest DMY (Figure 1a). As plot composition had no significant main effect, cultivars within the same ploidy level had similar DMY, and differences occurred mainly between diploid and tetraploid cultivars. Under regular cutting, tetraploid cultivars had significantly higher DMY than diploid cultivars (+7.01 dt DM ha^-1^ a^-1^, *P* < 0.01), whereas this difference was smaller and not significant under simulated grazing (+1.67 dt DM ha^-1^ a^-1^, *P* > 0.05; Supplementary Table S9). In contrast, under grazing, diploid cultivars tended to have higher DMY, although this was not significant (+3.72 dt DM ha^-1^ a^-1^, *P* > 0.05). This interaction between ploidy and defoliation treatment was significant between regular cutting and grazing (Supplementary Table S10). Cultivar rankings for DMY did not correlate significantly between any pair of defoliation treatments (Table 3).

**Fig. 1:**
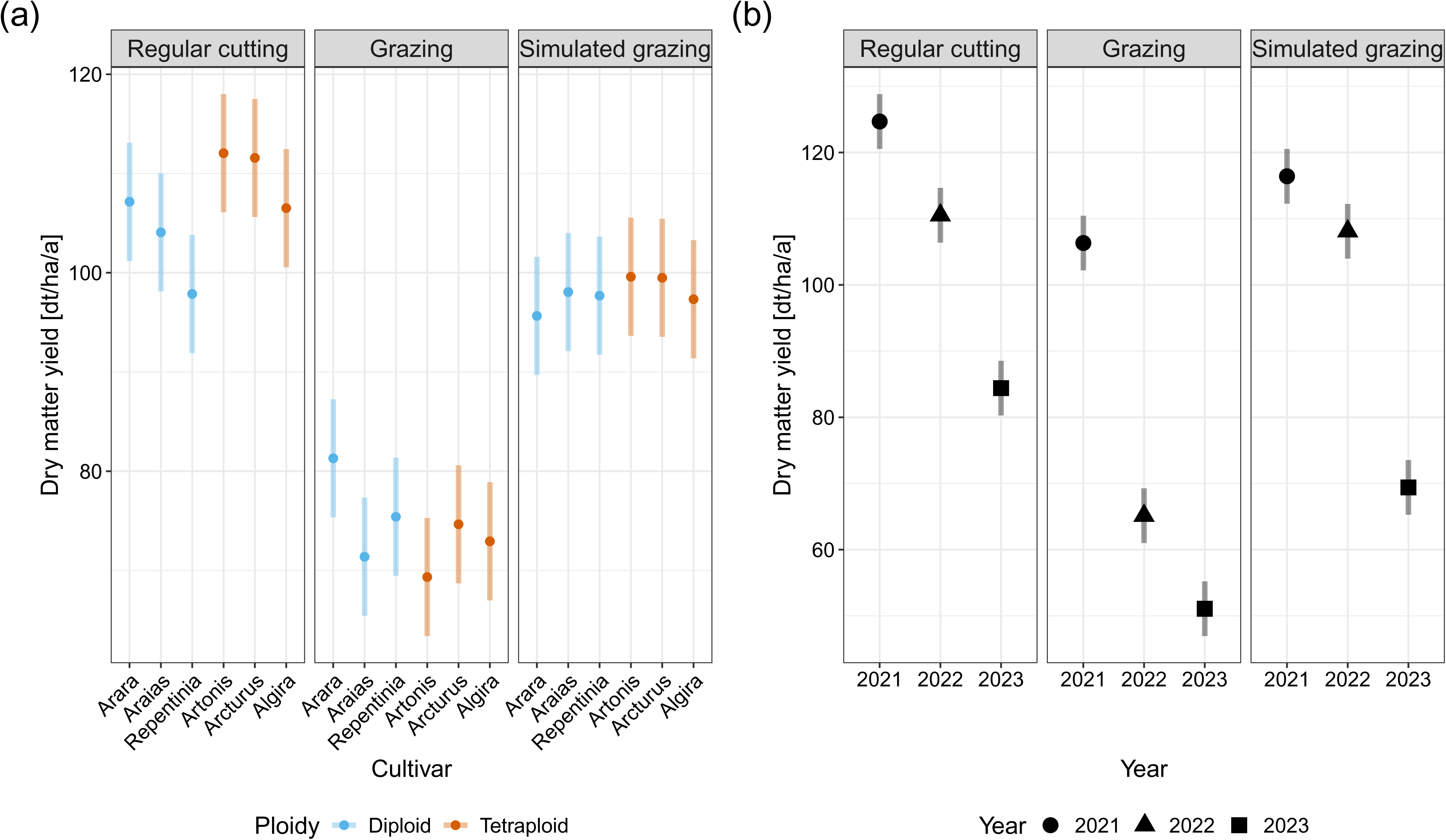
Estimated marginal means of the yearly sum of dry matter yield of six *Lolium perenne* L. cultivars per defoliation treatment (a) and summarised per defoliation treatment and year (b), bars indicate 95% confidence intervals.

**Table 2:** Analysis of variance of the yearly sum of dry matter yield of plot compositions based on six *Lolium perenne* L. cultivars, presenting the sum of squares (Sum Sq), the numerator and denominator degrees of freedom (df_Num_, df_Den_) of the *F*-test, the partial R² of each fixed effect, *F*-values, and the associated significance levels

| Factor | Sum Sq | df<sub>Num</sub> | df<sub>Den</sub> | Partial R<sup>2</sup> | *F*-value |
| --- | --- | --- | --- | --- | --- |
| **Main effects** |  |  |  |  |  |
| Plot composition | 1000.5 | 8 | 105.1 | 0.084 | 1.20<sup>ns</sup> |
| Defoliation treatment | 21142.8 | 2 | 10.4 | 0.948 | 101.78<sup>****</sup> |
| Location | 13650.1 | 1 | 4.0 | 0.971 | 131.49<sup>****</sup> |
| Year | 151300.0 | 2 | 240.0 | 0.859 | 728.71<sup>****</sup> |
| Residuals | 44743.7 | 431 |  |  |  |
| **Interactions** |  |  |  |  |  |
| Plot composition × defoliation treatment | 2767.6 | 13 | 105.1 | 0.202 | 2.05<sup>*</sup> |
| Plot composition × year | 957.1 | 16 | 240.0 | 0.037 | 0.58<sup>ns</sup> |
| Defoliation treatment × year | 13947.8 | 4 | 240.0 | 0.359 | 33.59<sup>****</sup> |
| Plot composition × defoliation treatment × year | 2010.3 | 26 | 240.0 | 0.075 | 0.74<sup>ns</sup> |

ns: $P > 0.05$ , \*: $P \leq 0.05$ , \*\*: $P \leq 0.01$ , \*\*\*: $P \leq 0.001$ , \*\*\*\*: $P \leq 0.0001$ . Partial R<sup>2</sup> values are only directly comparable within the same df<sub>Den</sub> range (e.g., location and defoliation treatment on ~4–10 df; plot-composition-related terms on ~105 df; year-related terms on ~240 df).

**Table 3:** Spearman rank correlation of six *Lolium perenne* L. cultivars between regular cutting (C), grazing (G), and simulated grazing (SG), per parameter (yearly sum of dry matter yield [DMY], neutral detergent fibre [NDF], acid detergent fibre [ADF], acid detergent lignin [ADL], crude protein [CP], non-fibre carbohydrates [NFC], and metabolisable energy [ME]). Higher values indicate a more similar ranking of cultivars between two defoliation treatments.

| Parameter | C vs G | C vs SG | G vs SG |
| --- | --- | --- | --- |
| DMY | -0.31<sup>ns</sup> | 0.49<sup>ns</sup> | -0.71<sup>ns</sup> |
| NDF | 0.89<sup>*</sup> | 1.00<sup>**</sup> | 0.89<sup>*</sup> |
| ADF | 0.94<sup>*</sup> | 1.00<sup>**</sup> | 0.94<sup>*</sup> |
| ADL | 0.94<sup>*</sup> | 1.00<sup>**</sup> | 0.94<sup>*</sup> |
| CP | 0.89<sup>*</sup> | 0.83<sup>ns</sup> | 0.89<sup>*</sup> |
| NFC | 0.77<sup>ns</sup> | 0.94<sup>*</sup> | 0.83<sup>ns</sup> |
| ME | 0.89<sup>*</sup> | 1.00<sup>**</sup> | 0.89<sup>*</sup> |

ns: $P > 0.05$ , \*: $P \leq 0.05$ , \*\*: $P \leq 0.01$

Over the years, DMY decreased, with a sharp decrease from 2021 to 2022 and from 2022 to 2023 in swards under grazing and simulated grazing, respectively (Figure 1b). The DMY of the mixtures in relation to the corresponding cultivars differed depending on the defoliation treatment. Under regular cutting, all mixtures generally had a DMY ranging between that of the corresponding cultivars (Supplementary Figure S3). Under grazing, the diploid mixture had a DMY ranging between the corresponding cultivars, while mixtures of a diploid and a tetraploid cultivar had a slightly higher DMY than the corresponding cultivars, although not significant (+4.49 dt DM ha^-1^ a^-1^, *P* > 0.05; Supplementary Table S11).

With increasing years, the similarity of the mixtures to Ari generally decreased under both regular cutting and grazing (Figure 2a). Under regular cutting, similarity to Ari decreased from 76% to 10% for Rep50 and from 63% to 47% for Alg50, but increased from 66% to 84% for Art50 (2021 to 2023). Under grazing, similarity to Ari decreased for all mixtures: from 81% to 33% for Rep50, from 85% to 45% for Art50, and from 84% to 73% for Alg50 (2021 to 2023).

**Fig. 2:**
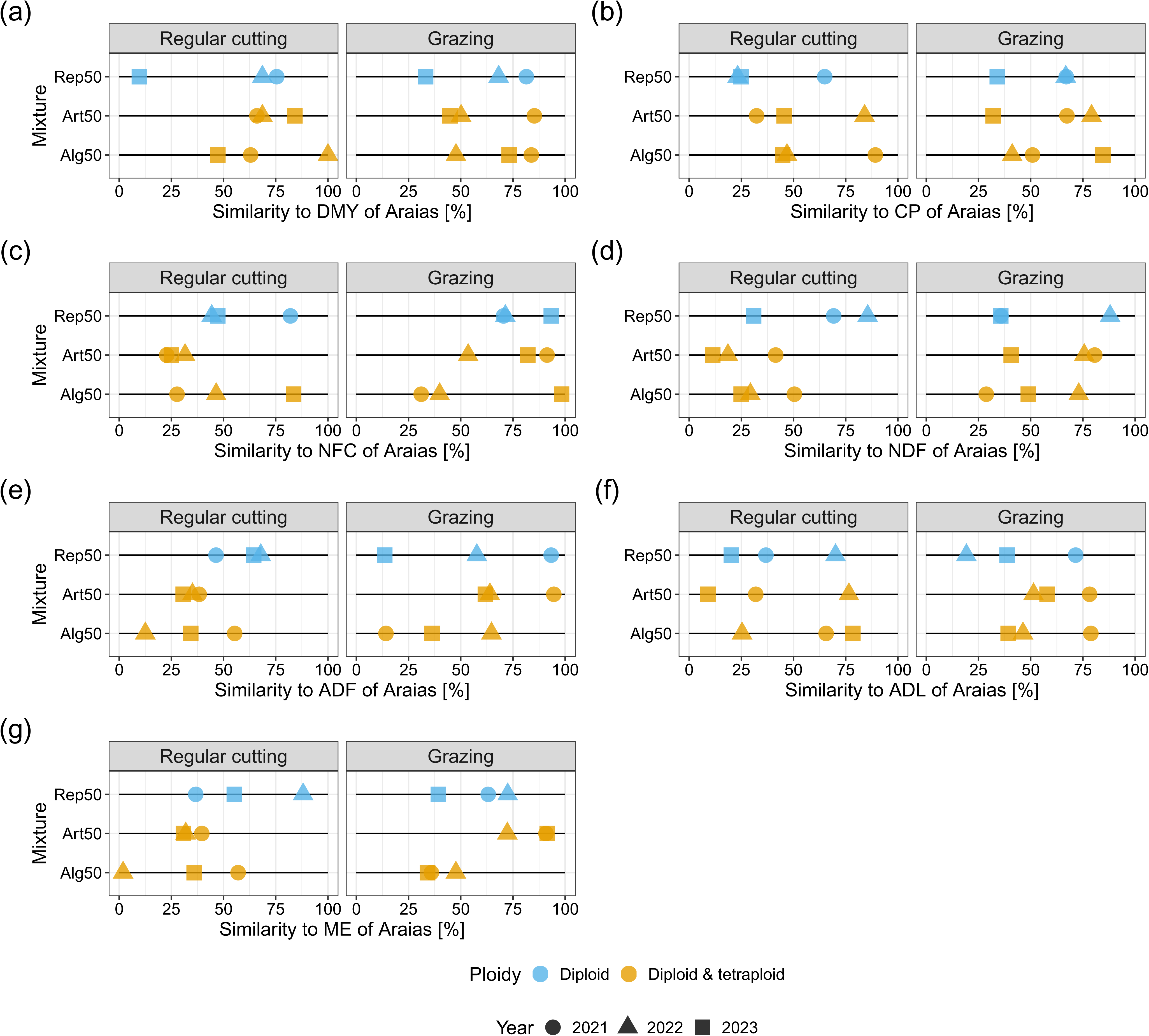
Relative similarity of estimated marginal means of mixtures to ‘Araias’ of yearly sum of dry matter yield (DMY; a), crude protein (CP; b), non-fibre carbohydrates (NFC; c), neutral detergent fibre (NDF; d), acid detergent fibre (ADF; e), acid detergent lignin (ADL; f), and metabolisable energy (ME; g) per defoliation treatment and year. Mixtures consisted of two cultivars of *Lolium perenne* L., with each mixture containing ‘Araias’ sown at a 50:50-ratio with ‘Repentinia’ (Rep50), ‘Artonis’ (Art50) or ‘Algira’ (Alg50). Mixtures with relative similarities of 100% or 0% have identical values to ‘Araias’ or the other cultivar, respectively.

### 3.2 Forage quality

Forage quality parameters were mainly affected by the defoliation, i.e. the sampled defoliation event within the year. As a main effect, its partial R^2^ ranged between 0.331 (NDF) and 0.682 (ADL), and its interaction with defoliation treatment between 0.049 (CP) and 0.348 (ME; Supplementary Table S7, Supplementary Table S8). Plot composition significantly affected all forage quality parameters (*P* < 0.0001), with partial R^2^ ranging between 0.020 (CP) and 0.101 (ME). Plot compositions under grazing and simulated grazing generally had similar values, whereas those under regular cutting were more distinct (Figure 3). Under regular cutting, NDF and ADF contents were higher, ADL and NFC contents were similar, and CP content and ME were lower compared to grazing and simulated grazing. As plot composition had only a minor effect on forage quality, differences occurred mainly between diploid and tetraploid cultivars: tetraploid cultivars were lower in structural components (i.e., NDF, ADF, and ADL) and similar or higher in cell contents (i.e., CP and NFC) and ME when compared to diploid cultivars (Supplementary Table S9). For most forage quality parameters, cultivar rankings were most similar between regular cutting and simulated grazing (Table 3).

**Fig. 3:**
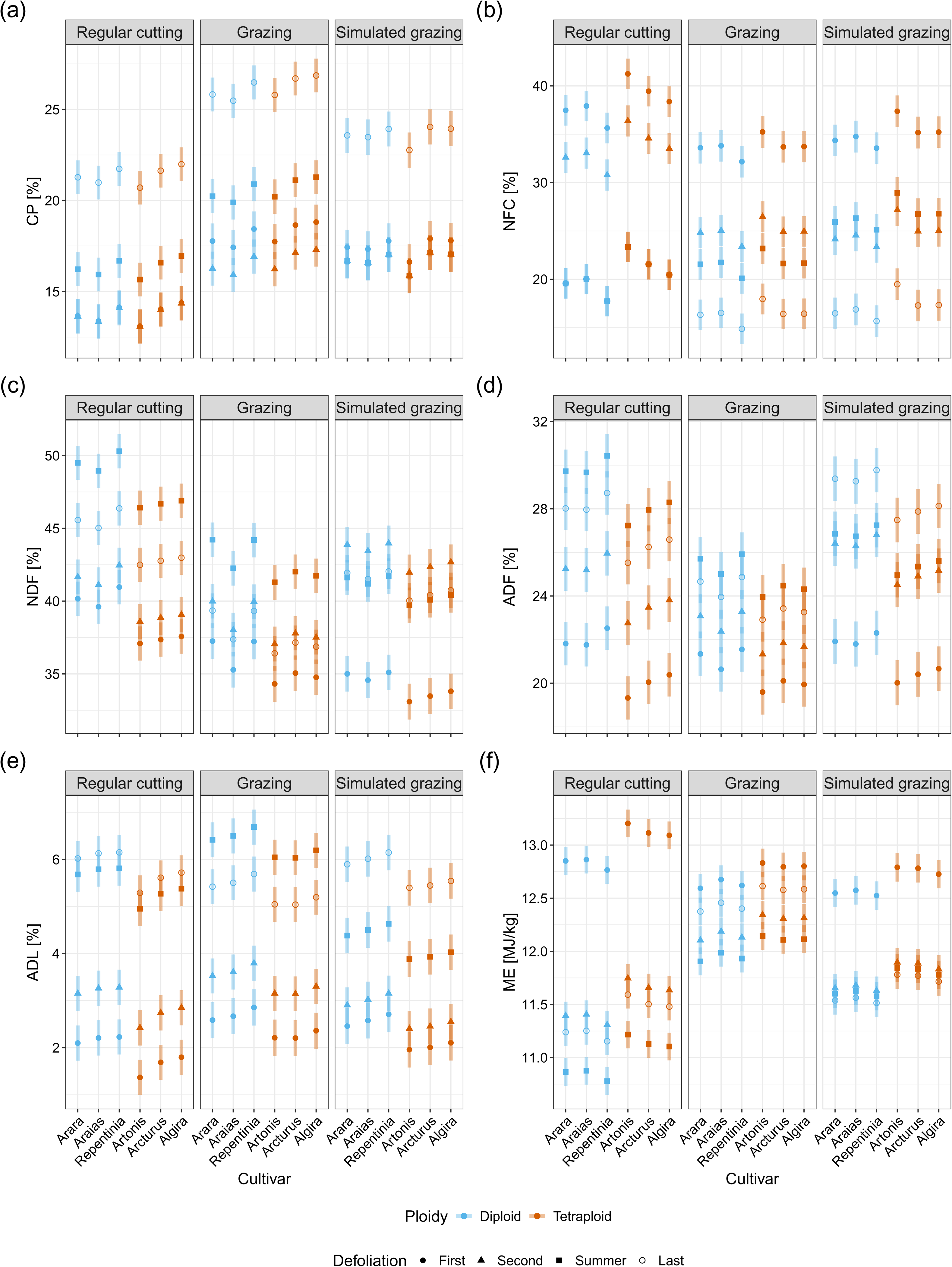
Estimated marginal means of crude protein (CP; a), non-fibre carbohydrates (NFC; b), neutral detergent fibre (NDF; c), acid detergent fibre (ADF; d), acid detergent lignin (ADL; e), and metabolisable energy (ME; f) of six *Lolium perenne* L. cultivars per defoliation treatment and defoliation, bars indicate 95% confidence intervals.

Forage quality values of the mixtures generally ranged between those of the corresponding cultivars (Figure 4; Supplementary Table S11). While mixtures of a diploid and a tetraploid cultivar had forage quality values closer to the tetraploid cultivar under regular cutting, they were closer to the diploid cultivar under grazing. From 2021 to 2023, the similarity of the mixtures to Ari decreased for most parameters (Figure 2b-g). By 2023, most mixtures tended to be less similar to Ari (i.e., similarity to Ari < 50%) in most forage quality parameters under both regular cutting and grazing. Under regular cutting, this was not the case for NFC, ADF, ADL, and ME, where few mixtures were more similar to Ari. Under grazing, the same was true for CP, ADF, ADL, and ME, whereas for NFC, all mixtures were more similar to Ari.

**Fig. 4:**
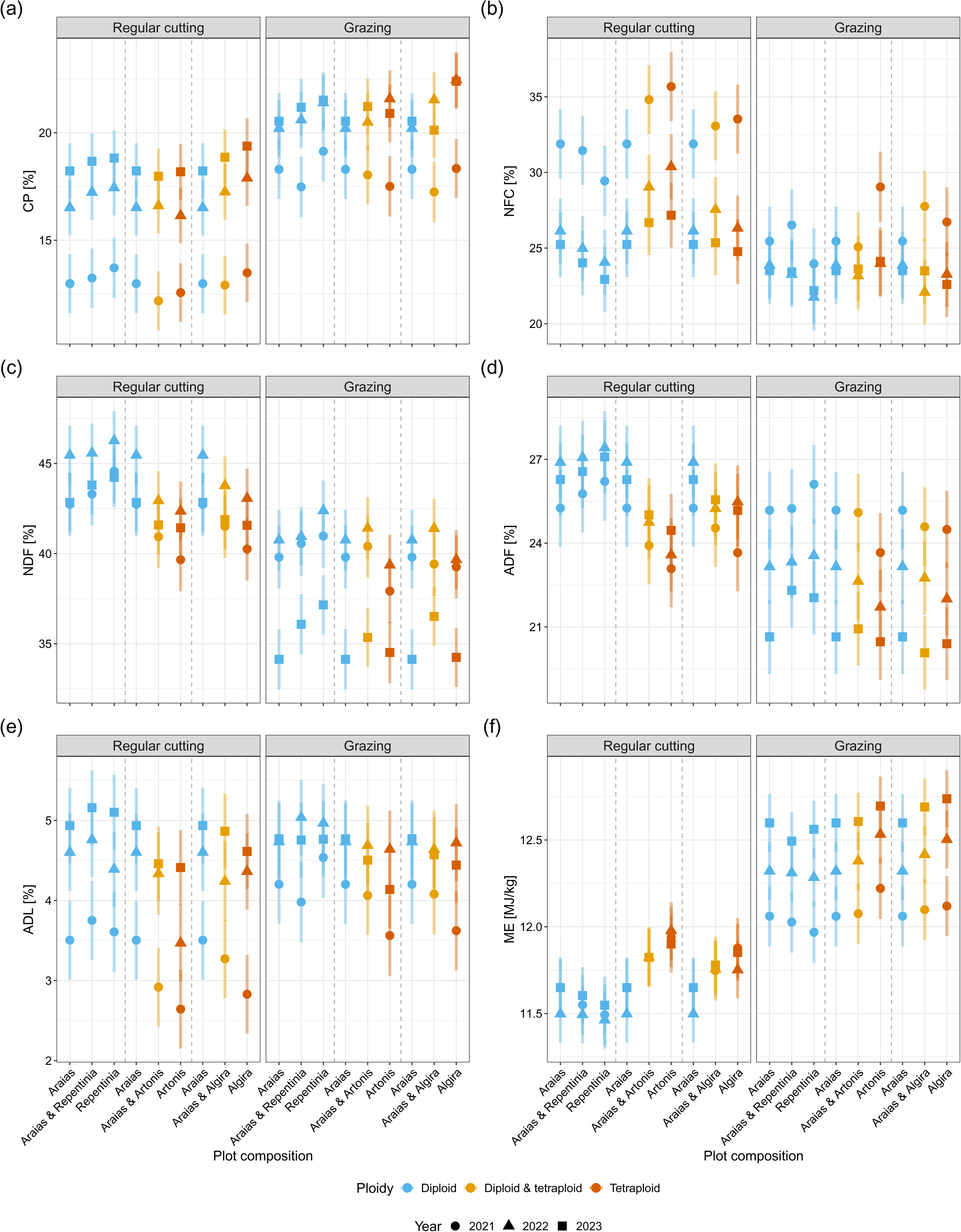
Estimated marginal means of crude protein (CP; a), non-fibre carbohydrates (NFC; b), neutral detergent fibre (NDF; c), acid detergent fibre (ADF; d), acid detergent lignin (ADL; e), and metabolisable energy (ME; f) per defoliation treatment, plot composition based on six *Lolium perenne* L. cultivars, and year, bars indicate 95% confidence intervals. Mixtures sown at a 50:50-ratio and the corresponding single-cultivar stands are shown in groups (separated by dashed lines) for direct comparison.

## 4 Discussion

By combining multi-year field data from three contrasting defoliation treatments at two sites, this study shows how defoliation treatment, ploidy, and cultivar mixing interact to determine the productivity and forage quality of perennial ryegrass swards. Five main findings emerged. First, productivity was determined primarily by year and defoliation treatment rather than by cultivar. Both the absolute values and the cultivar ranking differed between grazing and simulated grazing, suggesting that simulated grazing is not a suitable proxy for the DMY of grazed swards. Second, the effect of ploidy on DMY depended on the defoliation treatment and could be connected to differences in tiller densities. Third, forage quality was affected mainly by defoliation frequency and, to a lesser extent, by ploidy, and both the absolute values and the cultivar rankings were comparable between grazing and simulated grazing, making simulated grazing a suitable proxy for the forage quality of grazed swards. Fourth, cultivar mixtures generally were similar in DMY and forage quality compared to their components, with slightly higher DMY restricted to diploid-tetraploid mixtures under grazing. Fifth, the indirect inference of plot-composition shifts from sward performance (i.e., DMY and forage quality), a new approach, showed a similar pattern for most parameters and could become a valuable tool once validated against a reliable ground truth.

### 4.1 Year and defoliation treatment drive yield

Year and defoliation treatment were the main drivers of DMY, whereas plot composition had no significant effect. Across defoliation treatments, swards under regular cutting had the highest DMY, those under simulated grazing were intermediate, and grazed swards had the lowest DMY. This pattern is consistent with the general observation that less frequently defoliated swards produce higher DMY than more frequently defoliated swards (Binnie and Chestnutt, 1991; Boswell, 1977; Cashman et al., 2016; Wilkins, 1989). Because DMY and cultivar ranking differed between grazing and simulated grazing, simulated grazing was not a suitable proxy for the DMY of grazed swards. This may be explained by carry-over effects between rotations, which simulated grazing does not capture: cultivars grazed more heavily at a defoliation date enter the next rotation with less residual leaf area, which can carry over as reduced regrowth (Byrne et al., 2018; Tubritt et al., 2020a). Because cows could choose between plots, these differences in residual leaf area may have been larger than in a uniformly sown paddock. However, other studies have not revealed such DMY and ranking differences, possibly due to higher fertiliser input (Cashman et al., 2016; Creighton et al., 2012), higher precipitation (Creighton et al., 2012; Grace et al., 2019), and less intensive defoliation treatment with longer recovery phases between defoliations (Cashman et al., 2016; Creighton et al., 2012; Grace et al., 2019). Over the three years, DMY decreased in all defoliation treatments, with a large decrease in 2022 and 2023 for swards under grazing and simulated grazing, respectively. For the grazing plots, this decline could be attributed to the higher stress induced by grazing: cow treading, gripping and tearing of leaves, spatially heterogeneous excretion, and, due to high defoliation frequency, and thus, short recovery phases, more frequent periods of low sward height resulting in increased soil temperatures (Yan et al., 2018). The dry weather of 2022 and 2023, to which perennial ryegrass is poorly adapted, could have further decreased DMY and contributed to the year-to-year variability (Hearn et al., 2021; Tubritt et al., 2020b). Under simulated grazing, the decline occurred one year later than in the grazing plots, which could be attributed to the lower stress (i.e., less frequent defoliation, no treading, and no heterogeneous nutrient return) induced by mechanical harvesting.

### 4.2 Ploidy effects depend on defoliation treatment

The effect of ploidy on DMY depended on the defoliation treatment: Tetraploid cultivars had slightly higher DMY than diploid cultivars under regular cutting and simulated grazing, whereas diploid cultivars tended to have higher DMY than tetraploid cultivars under grazing. This is in accordance with other studies (Balocchi and López, 2009; Tubritt et al., 2020a, 2021) and can be explained by different tiller morphologies and densities. Diploid cultivars formed denser swards than tetraploid cultivars in our experiment (Käch et al., 2024), as in other studies (Byrne et al., 2018; Tubritt et al., 2020a, 2020b). They leave more residual leaf area, even at low post-grazing heights, and regrow more reliably under frequent grazing (Sills et al., 1982). Tetraploid cultivars had a lower tiller density (Käch et al., 2024) and are preferentially grazed by cattle, which lowers post-grazing height and further limits regrowth (Byrne et al., 2018; Tubritt et al., 2020a, 2020b, 2021). In the current study, cows could choose between plots of different composition, as in most plot trials. Where cows have no such choice, the difference in post-grazing height between ploidy levels may be smaller (Tubritt et al., 2020b), and with it the lower DMY of tetraploid cultivars compared to diploid cultivars. However, lower post-grazing height may not be the only reason, as diploid cultivars also produced more herbage when all plots were mown to the same height after each grazing (Balocchi and López, 2009). Grazing each plot composition as a separate paddock would allow testing this in future trials.

### 4.3 Forage quality reflects defoliation frequency and ploidy

Forage quality was affected mainly by defoliation frequency and, to some extent, by ploidy. Across forage quality parameters, defoliation was the dominant factor (partial R^2^ 0.331-0.682), because more frequent defoliation generally leads to younger plants at harvesting with higher leaf-to-stem ratio (Terry and Tilley, 1964). This results in a higher fraction of cell content (CP and NFC), a lower fraction of structural components (NDF, ADF, ADL), and thus a higher ME. Accordingly, regular cutting, the defoliation treatment with the lowest defoliation frequency, resulted in higher NDF and ADF, similar ADL, and lower CP and ME contents compared to grazing and simulated grazing. This is in agreement with the higher digestibility reported under more frequent defoliation (Bruinenberg et al., 2002; Cashman et al., 2016). Because plant maturity also depends on growing conditions, forage quality varied more within than between years: structural components were higher and cell contents lower in summer compared with the other time points, consistent with other seasonal analyses (Balocchi and López, 2009; Chesney et al., 2026; Colas et al., 2022). Besides these defoliation treatment and seasonal effects, tetraploid cultivars had similar or higher cell contents and ME and lower structural components than diploid cultivars, which can be attributed to the larger cells of tetraploid cultivars and correspondingly their higher volume-to-surface ratio (Sugiyama, 2005), and agrees with previous ploidy comparisons (Beecher et al., 2015; Byrne et al., 2018; Konkolewska et al., 2024; Tubritt et al., 2020a). Because plots under grazing and simulated grazing had comparable values across all parameters and cultivars ranked similarly, simulated grazing is a suitable proxy for forage quality under grazing, which has been reported before (Cashman et al., 2016). Furthermore, regular cutting ranked the cultivars similarly to grazing for most parameters, making it a suitable proxy for the ranking as well.

### 4.4 Performance of cultivar mixtures

Cultivar mixtures generally had performances ranging between those of their components. Forage quality values of all mixtures generally ranged between those of the corresponding cultivars across all parameters and defoliation treatments, consistent with other studies (Jones and Roberts, 1994; Tubritt et al., 2021), and DMY did so as well under regular cutting. Under grazing, however, diploid-tetraploid mixtures were similar or slightly higher in DMY than their components. This pattern is consistent with the observation that mixtures achieve higher DMY than their components when these differ morphologically (Rhodes, 1970): combining the many small tillers of diploid cultivars with the few large tillers of tetraploid cultivars allows the cultivars to compensate for one another (Beecher et al., 2015; Käch et al., 2024). That Rhodes (1970) reported even higher DMY advantage of mixtures over their components under regular cutting than under simulated grazing, using genotypes with strongly contrasting plant morphologies, is consistent with this interpretation, as more complementary mixtures tend to exceed the DMY of their components more (Trenbath, 1974). Because the cultivars used here differed less morphologically than those in Rhodes (1970), the resulting mixture effects were correspondingly smaller.

### 4.5 Inferring plot-composition shifts

The indirect inference of plot-composition shifts in mixtures, based on the similarity of mixtures to their components in DMY and forage quality, showed a similar pattern for most parameters. Similarity of the diploid-tetraploid and diploid-diploid mixtures generally decreased, suggesting a shift away from Ari. For diploid-tetraploid mixtures under grazing, this is despite the higher tiller density and grazing persistence of diploid cultivars (Käch et al., 2024; Sills et al., 1982) and the preferential grazing of tetraploid cultivars (Balocchi and López, 2009; Tubritt et al., 2021), which would rather suggest a growing dominance of the diploid cultivar. This could be explained by a general competitiveness of tetraploid cultivars, also reported under mechanical defoliation (Gilliland et al., 2011). Although the trend away from Ari was similar across most parameters, exceptions occurred, indicating that these results might not be sufficient to confirm compositional shifts. A suitable parameter would need to reflect the true shift in composition, without being confounded by phenotypic plasticity, whereby individual cultivars can express different phenotypes in mixture than in single-cultivar stands, independent of any actual shift in composition (Stefan et al., 2025). A reliable ground truth would help identify such a parameter. Direct methods – allozyme electrophoresis (Gilliland et al., 2011; Kennedy et al., 1985; Quaite and Camlin, 1986) or molecular genetic methods (van der Laat et al., 2025; Liu et al., 2010) – have allowed to estimate compositions with higher certainty but are labour-intensive. Recently developed approaches using genotyping-by-sequencing and amplicon sequencing require comparatively less labour and while remaining accurate could therefore reduce this barrier (Käch et al., 2026). Pairing such molecular methods with the performance-based approach used here would allow the identification of traits that estimate composition reliably. Once validated against such a ground truth, this approach could be valuable at several steps of cultivar development: shifts in composition reveal differences in persistence and competitiveness to breeders, and they help seed producers identify and farmers use cultivars that maintain their share and persist under grazing.

## 5 Conclusion

Productivity and forage quality were affected more strongly by defoliation treatment, year, and defoliation frequency than by the plot composition. Within this set of modern, recommended early-heading cultivars, cultivar choice therefore offers limited scope for improving sward performance compared with matching a cultivar to its defoliation treatment. The effect of ploidy depended on the defoliation treatment: tetraploid cultivars had slightly higher DMY under regular cutting and simulated grazing and similar or higher cell contents and metabolisable energy across defoliation treatments, whereas diploid cultivars had slightly higher DMY under grazing. Diploid cultivars may therefore be better suited to grazed swards and tetraploid cultivars where they are mown, although this yield-based recommendation is opposed by the higher forage quality of tetraploid cultivars, and the two criteria need to be weighed against each other. Mixtures of a diploid and a tetraploid cultivar had slightly higher DMY than their components under grazing and forage quality was intermediate to their components under grazing and regular cutting. Such mixtures may therefore be suitable for swards used for both mowing and grazing, or where the future use is not yet determined. Simulated grazing reproduced the forage quality of grazed swards, both in absolute values and cultivar ranking, but did not reproduce the cultivar ranking for DMY. Forage quality can therefore be assessed under mechanical defoliation, whereas yield rankings intended to inform grazing require evaluation under grazing. The inference of plot-composition shifts from sward performance was similar across traits; validating this approach against marker-based methods would establish a ground truth and make compositional monitoring feasible at the scale of variety testing. Evaluating cultivars and mixtures under grazing, with each plot composition grazed as a separate paddock to exclude selective grazing, would narrow the gap between how cultivars are tested and how they are used in practice.

## 6 Acknowledgements

We thank Hansueli Hirschi (Agroscope) for sowing and maintaining the field experiments and for harvesting the plots under regular cutting and simulated grazing. We thank Hansjürg Fuhrimann, Hugo Jung, and Fabio Tanner for providing space to conduct the field experiments and coordinating the defoliations by their cows. We thank Simon Ineichen (BFH-HAFL) for scientific advice and administrative support. We thank Tamara Köke (BFH-HAFL), Michael Howald (BFH-HAFL), Simon Jäggi (BFH-HAFL), Elias Marbach (BFH-HAFL), Oliver Stettler (BFH-HAFL), and Micha Wyss (BFH-HAFL) for their support on conducting measurements in the field experiments, milling samples, and/or measuring forage quality. We thank Martina Müller (BFH-HAFL) for her expertise and advice regarding NIRS measurements and calibration construction. We thank InnoSuisse PSI, Swiss Grassland Society (AGFF), Eric Schweizer Ltd, Fenaco CP UFA-Samen, Otto Hauenstein Seeds Inc., Samen Steffen Ltd, and Delley seeds and plants Ltd for funding this project.

## Supporting Information A Materials and Methods

### A.1 Excretion estimation

Excretion by cows during grazing was estimated using the following formula:

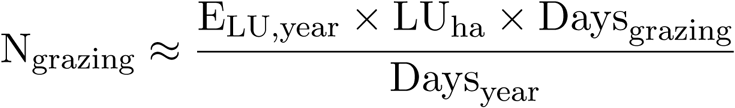

where N_grazing_ denotes the estimated nitrogen excreted on grazing plots per hectare and year, and E denotes the estimated excretion of N per livestock unit (LU) and year (Richner and Sinaj, 2017, Table 2, p. 60). For each of the nine grazing events, the cows grazed for two consecutive half days. This resulted in the following:

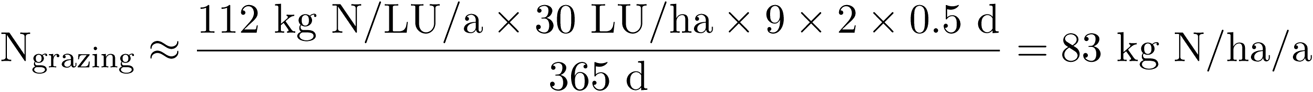

### A.2 Soil samples

Soil samples were collected using the Swiss Sampler (Eric Schweizer, Thun, Switzerland) and analysed by Eric Schweizer (Thun, Switzerland). For each location, samples were collected at the start of the experiment (October 2019) and during the experiment (March 2022). At the first and second time point, nine and 15 samples were collected at each location, respectively (Figure S2). More samples were taken at the second time point to account for potential effects of defoliation treatment. To have one representative sample per area, ten subsamples were collected, mixed, and pooled.

### A.3 Forage quality calibration

Forage quality was analysed by measuring samples of the first, second, summer, and last defoliation. Samples of the summer defoliation were collected between mid-June and mid-July, depending on the defoliation time point of each defoliation treatment. Forage quality was assessed using a near-infrared (NIR) spectrometer NIRFlex N-500 (Büchi, Flawil, Switzerland). For this, 168 representative samples, evenly distributed across years, defoliations, locations, and cultivars, were analysed using wet-chemistry by Dairy One (Ithaca, NY, USA). NIR spectra from these measurements were acquired in triplicates (resulting in 504 measurements) and used as references to develop quantitative calibrations using the NIRCal software suite (Büchi, Flawil, Switzerland; Table S6).

### A.4 Forage quality equations

Digestibility of *Lolium perenne* L. in pure swards (Agroscope, 2021):

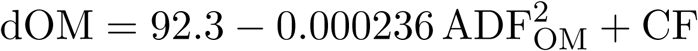

where dOM is the digestibility of organic matter (in %), ADF_OM_ is the organic part of acid detergent fibre (ADF, in g/kg organic matter), and CF is the correction factor added to or subtracted from dOM based on regrowth and development stage (Table S1). For grazing, simulated grazing, and regular cutting, the correction factors of DS 2, DS 2, and DS 3 were used, respectively. ADF_OM_ was approximated by

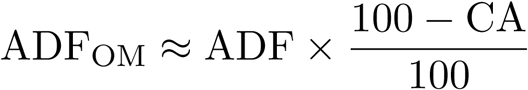

where CA is the crude ash (in %).

**Table S1:** Correction factor added to or subtracted from calculated digestibility of organic matter based on defoliation number of the year and development stage (DS) of *Lolium perenne* L. DS 1 to 5 correspond to tillering, stem elongation, start of heading, full heading, and end of heading, respectively (adapted from Agroscope (2021)).

|  | DS 1 | DS 2 | DS 3 | DS 4 | DS 5 |
| --- | --- | --- | --- | --- | --- |
| 1<sup>st</sup> defoliation | -2 | 0 | 3 | 5 | 4 |
| 2<sup>nd</sup> and following defoliations | 1 | 0 | 0 | -3 | -2 |

For calculating ME, dOM was converted to g/kg and used in the following equation (Agroscope, 2021):

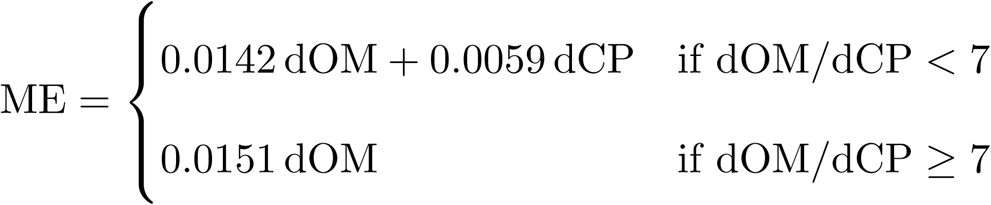

where dCP is the digestible crude protein (in g/kg):

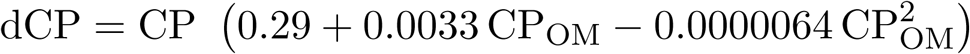

where CP is the crude protein (in g/kg) and CP_OM_ is the organic part of CP (in g/kg organic matter) which was approximated by

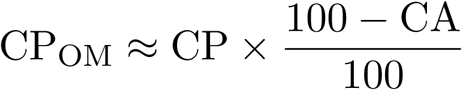

### A.5 Data analysis

Data processing, analysis, and visualisation was conducted with R v4.5.0 (R core team, 2025) in RStudio v2025.09.2.418 (Posit team, 2025) using readxl v1.4.5 (Wickham and Bryan, 2025), dplyr v1.1.4 (Wickham et al., 2022), tidyr v1.3.1 (Wickham and Girlich, 2022), stringr v1.5.1 (Wickham, 2022), lubridate v1.9.4 (Grolemund and Wickham, 2011), purrr v1.0.4 (Wickham and Henry, 2023), randomForest v4.7.1.2 (Liaw and Wiener, 2002), e1071 v1.7.16 (Meyer et al., 2024), caret v7.0.1 (Kuhn, 2008), lmerTest v3.1.3 (Kuznetsova et al., 2017), pbkrtest v0.5.4 (Halekoh and Højsgaard, 2014), r2glmm v0.1.3 (Jaeger, 2025), emmeans v1.11.1 (Lenth, 2025), sjPlot v2.8.17 (Lüdecke, 2024), ggplot2 3.5.2 (Wickham, 2009), ggpubr v0.6.0 (Kassambara, 2022), scales v1.4.0 (Wickham et al., 2025), patchwork v1.3.0 (Pedersen, 2024), quarto v1.7.32 (Allaire et al., 2025), shiny v1.10.0 (Chang et al., 2024), kableExtra v1.4.0 (Zhu, 2024), grid v4.5.0 (R core team, 2025), magick v2.9.0 (Ooms, 2025a), pdftools v3.6.0 (Ooms, 2025b), glue v1.8.0 (Hester and Bryan, 2024), and conflicted v1.2.0 (Wickham, 2023).

## Supporting Information B Figures and tables

**Table S2:** Dates of defoliation (mm-dd) of *Lolium perenne* L. swards and sampling for forage quality measurements in Langenthal BE (LT) and Eschenbach LU (EB) in the sampling years 2021, 2022, and 2023.

| Year | Defoliation | Regular cutting | Grazing |  | Simulated grazing |  |
| --- | --- | --- | --- | --- | --- | --- |
|  |  |  | LT | EB | LT | EB |
| 2021 | 1 | 04-28<sup>†‡</sup> | <sup>‡</sup> | 05-05<sup>†</sup> | 04-14<sup>†‡</sup> | 04-14<sup>†</sup> |
|  | 2 | 06-02<sup>†‡</sup> | 06-14<sup>†</sup> | 05-26<sup>†</sup> | 05-12<sup>†‡</sup> | 05-12<sup>†</sup> |
|  | 3 | 07-09<sup>†</sup> | 07-14<sup>†</sup> | 06-18 | 06-10 | 06-10 |
|  | 4 | 08-26 | 08-06 | 07-07<sup>†</sup> | 07-09<sup>†</sup> | 07-09<sup>†</sup> |
|  | 5 | 10-11<sup>†</sup> | 08-27 | 07-29 | 08-05 | 08-05 |
|  | 6 |  | 09-21 | 08-17 | 08-26 | 08-26 |
|  | 7 |  | 10-30<sup>†</sup> | 09-09 | 10-11 | 09-30 |
|  | 8 |  |  | 10-04 | 11-05<sup>†</sup> | 11-05<sup>†</sup> |
|  | 9 |  |  | 11-08<sup>†</sup> |  |  |
| 2022 | 1 | 05-02<sup>†</sup> | 04-22<sup>†</sup> | 04-21<sup>†</sup> | 04-11<sup>†</sup> | 04-11<sup>†</sup> |
|  | 2 | 06-03<sup>†</sup> | 05-20<sup>†</sup> | 05-14<sup>†</sup> | 05-09<sup>†</sup> | 05-09<sup>†</sup> |
|  | 3 | 07-18<sup>†</sup> | 06-09 | 06-04 | 06-03 | 06-03 |
|  | 4 | 08-29 | 07-02<sup>†</sup> | 07-01<sup>†</sup> | 07-04<sup>†</sup> | 07-04<sup>†</sup> |
|  | 5 | 10-06<sup>†</sup> | 07-28 | 07-22 | 08-04 | 08-04 |
|  | 6 |  | 08-19 | 08-12 | 08-29 | 08-29 |
|  | 7 |  | 09-08 | 09-04 | 10-06 | 10-06 |
|  | 8 |  | 10-01 | 09-30 | 11-02<sup>†</sup> | 11-02<sup>†</sup> |
|  | 9 |  | 10-29<sup>†</sup> | 10-28<sup>†</sup> |  |  |
| 2023 | 1 | 04-24<sup>†</sup> | 04-21<sup>†</sup> | 04-19<sup>†</sup> | 04-11<sup>†</sup> | 04-11<sup>†</sup> |
|  | 2 | 05-30<sup>†</sup> | 05-12<sup>†</sup> | 05-10<sup>†</sup> | 05-08<sup>†</sup> | 05-08<sup>†</sup> |
|  | 3 | 07-10<sup>†</sup> | 06-07 | 06-06 | 06-05<sup>†</sup> | 06-05<sup>†</sup> |
|  | 4 | 08-29 | 07-06<sup>†</sup> | 07-04<sup>†</sup> | 07-03 | 07-03 |
|  | 5 | 10-12<sup>†</sup> | 08-03 | 08-02 | 07-31 | 07-31 |
|  | 6 |  | 08-23 | 08-22 | 08-29<sup>†</sup> | 08-29<sup>†</sup> |
|  | 7 |  | 09-20 | 10-04 | 10-02 | 10-02 |
|  | 8 |  | 10-09 | 10-30<sup>†</sup> | 10-27 | 10-27 |
|  | 9 |  | 11-01<sup>†</sup> |  |  |  |

<sup>†</sup> Forage quality measurement
<sup>‡</sup> Containing missing values

**Fig. S1:**
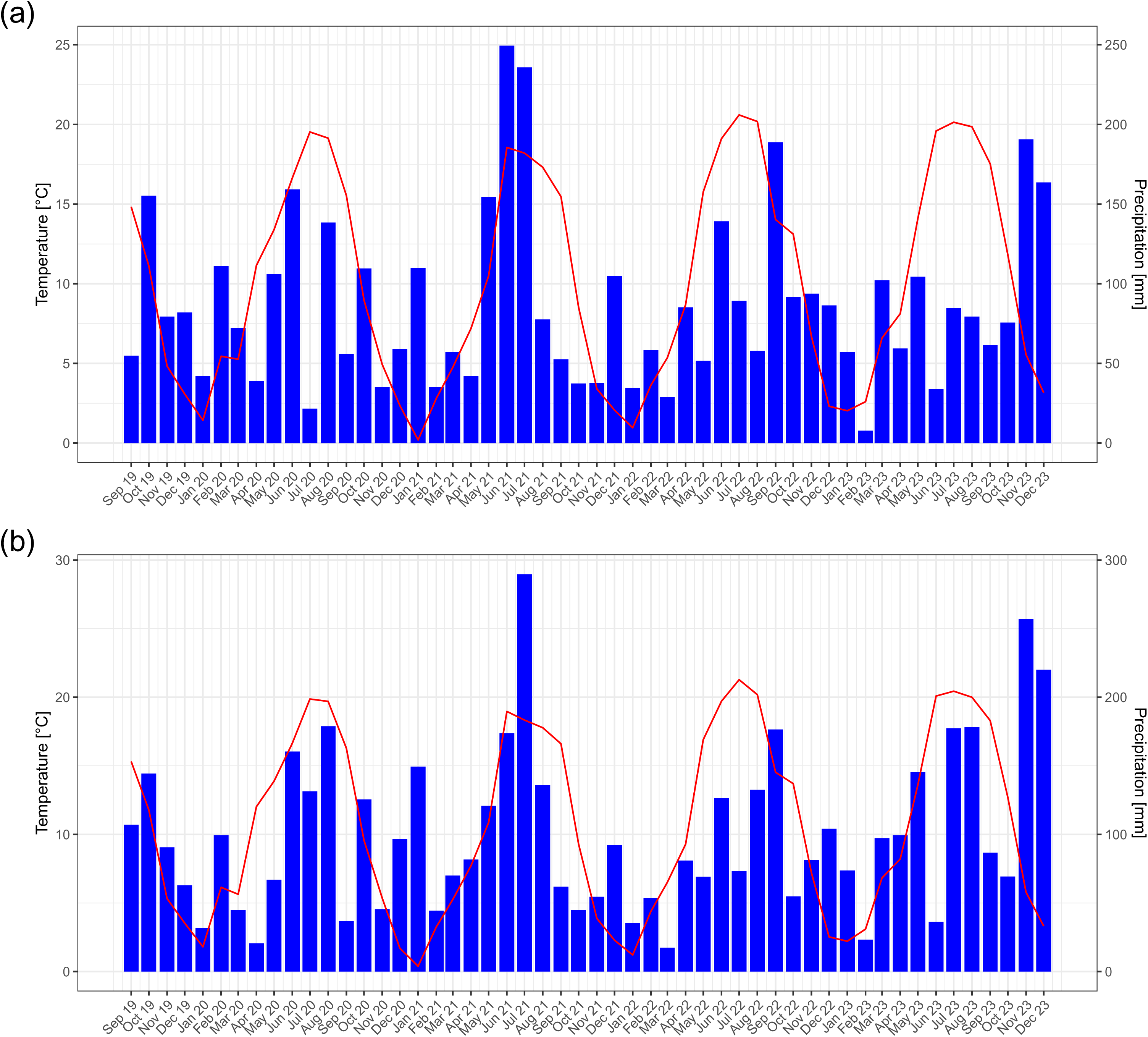
Temperature (monthly average) and precipitation (monthly sum) in Langenthal BE (a) and Eschenbach LU (b) from 2019 to 2023.

**Table S3:** Temperature (monthly average), precipitation (monthly sum), and relative humidity (monthly average) in Langenthal BE and Eschenbach LU from 2019 to 2023.

| Location | Parameter | Year | Jan | Feb | Mar | Apr | May | Jun | Jul | Aug | Sep | Oct | Nov | Dec | Total |
| --- | --- | --- | --- | --- | --- | --- | --- | --- | --- | --- | --- | --- | --- | --- | --- |
| Langenthal | Temperature [°C] | 2019 |  |  |  |  |  |  |  |  | 14.83 | 11.12 | 4.84 | 3.08 |  |
|  |  | 2020 | 1.43 | 5.45 | 5.26 | 11.15 | 13.39 | 16.63 | 19.53 | 19.14 | 15.54 | 8.97 | 4.94 | 2.36 | 10.32 |
|  |  | 2021 | 0.20 | 2.81 | 4.75 | 7.20 | 10.50 | 18.55 | 18.20 | 17.31 | 15.47 | 8.52 | 3.41 | 2.07 | 9.08 |
|  |  | 2022 | 0.96 | 3.64 | 5.36 | 8.69 | 15.77 | 19.11 | 20.60 | 20.18 | 14.03 | 13.12 | 6.69 | 2.31 | 10.87 |
|  |  | 2023 | 2.03 | 2.60 | 6.59 | 8.12 | 14.10 | 19.59 | 20.14 | 19.85 | 17.53 | 11.82 | 5.55 | 3.16 | 10.92 |
|  | Precipitation [mm] | 2019 |  |  |  |  |  |  |  |  | 54.80 | 155.20 | 79.40 | 82.00 |  |
|  |  | 2020 | 42.20 | 111.20 | 72.40 | 39.00 | 106.20 | 159.20 | 21.60 | 138.40 | 56.00 | 109.60 | 35.00 | 59.20 | 950.00 |
|  |  | 2021 | 109.80 | 35.20 | 57.20 | 42.20 | 154.60 | 249.40 | 235.80 | 77.60 | 52.60 | 37.40 | 37.80 | 104.80 | 1194.40 |
|  |  | 2022 | 34.60 | 58.40 | 28.80 | 85.20 | 51.60 | 139.20 | 89.20 | 57.80 | 188.80 | 91.70 | 93.80 | 86.40 | 1005.50 |
|  |  | 2023 | 57.20 | 7.80 | 102.20 | 59.40 | 104.40 | 34.00 | 84.80 | 79.40 | 61.40 | 75.60 | 190.60 | 163.60 | 1020.40 |
|  | Relative humidity [%] | 2019 |  |  |  |  |  |  |  |  | 80.50 | 90.65 | 91.79 | 91.15 |  |
|  |  | 2020 | 90.67 | 80.53 | 76.85 | 67.55 | 75.43 | 78.20 | 71.98 | 78.56 | 82.56 | 89.83 | 92.35 | 91.70 | 81.35 |
|  |  | 2021 | 88.63 | 84.30 | 76.39 | 67.68 | 76.50 | 76.98 | 81.04 | 79.53 | 82.14 | 85.24 | 88.62 | 89.02 | 81.34 |
|  |  | 2022 | 84.76 | 79.24 | 70.30 | 73.66 | 73.23 | 74.74 | 68.10 | 70.29 | 80.84 | 87.56 | 88.48 | 89.44 | 78.39 |
|  |  | 2023 | 85.31 | 80.54 | 78.24 | 78.02 | 76.50 | 65.64 | 72.67 | 74.97 | 78.61 | 83.08 | 85.95 | 87.16 | 78.89 |
| Eschenbach | Temperature [°C] | 2019 |  |  |  |  |  |  |  |  | 15.32 | 11.79 | 5.32 | 3.53 |  |
|  |  | 2020 | 1.80 | 6.14 | 5.63 | 12.02 | 13.88 | 16.64 | 19.87 | 19.70 | 16.28 | 9.57 | 5.35 | 1.70 | 10.72 |
|  |  | 2021 | 0.39 | 3.26 | 5.25 | 7.72 | 10.87 | 18.96 | 18.33 | 17.78 | 16.61 | 9.29 | 3.88 | 2.29 | 9.55 |
|  |  | 2022 | 1.21 | 4.40 | 6.51 | 9.29 | 16.91 | 19.71 | 21.28 | 20.19 | 14.53 | 13.70 | 7.20 | 2.53 | 11.45 |
|  |  | 2023 | 2.21 | 3.10 | 6.83 | 8.20 | 13.61 | 20.09 | 20.44 | 20.00 | 18.30 | 12.70 | 5.78 | 3.31 | 11.21 |
|  | Precipitation [mm] | 2019 |  |  |  |  |  |  |  |  | 107.10 | 144.30 | 90.60 | 62.90 |  |
|  |  | 2020 | 31.60 | 99.30 | 44.90 | 20.60 | 66.90 | 160.40 | 131.40 | 178.90 | 36.70 | 125.50 | 45.50 | 96.50 | 1038.20 |
|  |  | 2021 | 149.40 | 44.40 | 70.00 | 81.70 | 120.80 | 173.80 | 289.70 | 135.80 | 61.80 | 44.90 | 54.50 | 92.10 | 1318.90 |
|  |  | 2022 | 35.40 | 53.60 | 17.40 | 80.90 | 69.10 | 126.60 | 73.10 | 132.50 | 176.50 | 54.80 | 81.20 | 104.10 | 1005.20 |
|  |  | 2023 | 73.70 | 23.30 | 97.30 | 99.30 | 145.20 | 36.20 | 177.40 | 178.30 | 86.60 | 69.30 | 256.90 | 220.00 | 1463.50 |
|  | Relative humidity [%] | 2019 |  |  |  |  |  |  |  |  | 79.94 | 89.41 | 91.24 | 88.82 |  |
|  |  | 2020 | 89.50 | 72.88 | 75.33 | 63.31 | 72.59 | 77.90 | 71.84 | 77.08 | 82.00 | 88.98 | 93.53 | 95.78 | 80.06 |
|  |  | 2021 | 89.97 | 85.58 | 74.77 | 67.08 | 76.03 | 78.66 | 85.40 | 82.57 | 86.01 | 89.41 | 94.20 | 94.55 | 83.69 |
|  |  | 2022 | 89.27 | 78.55 | 66.17 | 73.66 | 74.71 | 77.16 | 65.94 | 72.97 | 81.07 | 89.92 | 91.33 | 94.13 | 79.58 |
|  |  | 2023 | 86.37 | 81.57 | 77.08 | 79.29 | 81.33 | 62.62 | 73.40 | 76.13 | 81.42 | 84.86 | 88.87 | 91.27 | 80.35 |

**Table S4:** Manure content of samples taken before and after winter. PH, dry matter (DM), organic matter and carbon (OM, C_org_), N, and macronutrients (P, K, Ca, Mg) were analysed by a direct measurement, drying at 105°C, loss on ignition method, Kjeldahl method, and ICP-OES measurement of microwave-assisted aqua regia extract, respectively.

| Location | Date | pH | DM [%] | OM [% DM] | C<sub>org</sub> [g/kg DM] | N [g/kg DM] | C/N | P [kg/t DM] | K [kg/t DM] | Ca [kg/t DM] | Mg [kg/t DM] |
| --- | --- | --- | --- | --- | --- | --- | --- | --- | --- | --- | --- |
| Langenthal | Mar 2020 | 7.23 | 3.8 | 72.9 | 422.6 | 43.7 | 9.67 | 7.80 | 65.5 | 26.2 | 5.25 |
|  | Oct 2020 | 7.33 | 1.6 | 61.8 | 358.1 | 30.8 | 11.62 | 7.86 | 36.6 | 48.5 | 6.30 |
|  | Apr 2021 | 8.15 | 10.2 | 63.6 | 368.9 | 44.4 | 8.32 | 6.09 | 79.8 | 32.9 | 6.17 |
|  | Mar 2022 | 7.23 | 6.8 | 75.4 | 437.3 | 33.8 | 12.93 | 7.83 | 41.5 | 39.8 | 5.34 |
|  | Nov 2022 | 7.04 | 6.1 | 66.0 | 382.8 | 26.4 | 14.49 | 7.47 | 14.6 | 44.1 | 3.75 |
|  | Mar 2023 | 7.66 | 6.2 | 75.1 | 435.2 | 40.0 | 10.88 | 6.62 | 59.8 | 26.5 | 5.10 |
| Eschenbach | Mar 2020 | 6.99 | 3.9 | 69.3 | 401.9 | 37.2 | 10.80 | 8.24 | 52.4 | 16.9 | 4.89 |
|  | Oct 2020 | 7.33 | 1.7 | 57.9 | 335.9 | 42.7 | 7.88 | 8.17 | 97.1 | 24.7 | 7.93 |
|  | Apr 2021 | 7.24 | 5.2 | 73.4 | 425.8 | 27.1 | 15.69 | 5.64 | 44.9 | 19.6 | 5.00 |
|  | Mar 2022 | 7.18 | 3.2 | 78.3 | 453.6 | 45.6 | 9.95 | 19.60 | 29.5 | 22.7 | 18.10 |
|  | Nov 2022 | 7.16 | 2.1 | 63.5 | 368.2 | 55.4 | 6.65 | 8.28 | 88.6 | 25.8 | 5.19 |
|  | Mar 2023 | 7.57 | 4.8 | 76.6 | 443.8 | 32.2 | 13.79 | 5.41 | 56.8 | 15.4 | 4.04 |

**Table S5:** Soil characteristics and content of samples taken at the beginning (2019) and during (2022) the experiment. PH, Soil texture (humus, clay, silt), and macronutrients (P, K, Mg) were analysed by pH (1:2.5 H_2_O), texture by feel method, and extraction with ammonium acetate EDTA (AAE10), respectively. Plot numbers are based on Figure S2.

| Location | Year | Plot | pH | Humus<br>[% w/w] | Clay<br>[% w/w] | Silt<br>[% w/w] | P<br>[mg/kg] | K<br>[mg/kg] | Mg<br>[mg/kg] |
| --- | --- | --- | --- | --- | --- | --- | --- | --- | --- |
| Langenthal | 2019 | 1 | 6.2 | 2.0 | 16 | 11 | 151.8 | 278.6 | 151.0 |
|  |  | 2 | 6.2 | 2.0 | 21 | 21 | 133.1 | 289.8 | 143.6 |
|  |  | 3 | 6.2 | 2.0 | 21 | 21 | 150.6 | 326.9 | 132.2 |
|  |  | 4 | 6.2 | 2.5 | 16 | 11 | 96.0 | 281.7 | 152.7 |
|  |  | 5 | 5.9 | 2.5 | 21 | 21 | 85.0 | 241.5 | 136.0 |
|  |  | 6 | 6.2 | 2.5 | 21 | 21 | 100.4 | 221.5 | 125.0 |
|  |  | 7 | 6.1 | 2.0 | 21 | 21 | 87.6 | 210.6 | 133.7 |
|  |  | 8 | 6.0 | 2.5 | 16 | 21 | 77.4 | 176.3 | 131.1 |
|  |  | 9 | 6.1 | 2.5 | 16 | 21 | 88.7 | 190.3 | 117.8 |
|  | 2022 | 1 | 6.1 | 1.0 | 11 | 21 | 145.2 | 205.5 | 160.9 |
|  |  | 2 | 6.0 | 1.5 | 11 | 11 | 114.0 | 181.4 | 146.0 |
|  |  | 3 | 6.1 | 0.0 | 11 | 11 | 146.5 | 257.8 | 160.6 |
|  |  | 4 | 6.3 | 2.5 | 11 | 11 | 166.7 | 282.9 | 155.1 |
|  |  | 5 | 5.8 | 2.5 | 11 | 11 | 135.7 | 258.3 | 150.2 |
|  |  | 6 | 6.6 | 2.5 | 11 | 11 | 166.6 | 326.4 | 146.4 |
|  |  | 7 | 6.3 | 2.5 | 11 | 11 | 115.4 | 374.3 | 162.6 |
|  |  | 8 | 5.9 | 2.5 | 11 | 11 | 84.0 | 234.7 | 133.1 |
|  |  | 9 | 6.1 | 3.0 | 11 | 11 | 116.3 | 378.0 | 139.2 |
|  |  | 10 | 5.9 | 2.5 | 11 | 11 | 83.4 | 176.0 | 153.7 |
|  |  | 11 | 5.8 | 2.5 | 11 | 11 | 74.6 | 237.2 | 158.8 |
|  |  | 12 | 5.7 | 2.5 | 11 | 11 | 77.7 | 172.4 | 135.6 |
|  |  | 13 | 6.0 | 2.0 | 11 | 11 | 101.4 | 279.5 | 131.4 |
|  |  | 14 | 6.0 | 2.5 | 11 | 11 | 91.5 | 280.7 | 140.4 |
|  |  | 15 | 5.9 | 2.5 | 11 | 11 | 125.4 | 348.2 | 126.4 |
| Eschenbach | 2019 | 1 | 6.8 | 3.0 | 26 | 21 | 176.6 | 118.1 | 178.9 |
|  |  | 2 | 7.1 | 2.5 | 21 | 21 | 123.4 | 121.6 | 166.9 |
|  |  | 3 | 7.2 | 3.0 | 21 | 31 | 128.1 | 139.2 | 184.1 |
|  |  | 4 | 6.9 | 2.5 | 21 | 21 | 142.2 | 155.0 | 190.4 |
|  |  | 5 | 7.0 | 2.5 | 26 | 21 | 82.3 | 105.2 | 160.5 |
|  |  | 6 | 6.9 | 2.5 | 21 | 21 | 95.6 | 113.2 | 150.9 |
|  |  | 7 | 7.0 | 2.5 | 21 | 31 | 160.4 | 158.4 | 187.5 |
|  |  | 8 | 7.1 | 2.5 | 26 | 21 | 117.3 | 137.1 | 173.1 |
|  |  | 9 | 7.1 | 2.5 | 21 | 21 | 153.3 | 105.3 | 165.8 |
|  | 2022 | 1 | 6.7 | 4.0 | 11 | 41 | 190.5 | 281.5 | 221.4 |
|  |  | 2 | 6.8 | 3.0 | 16 | 31 | 156.7 | 227.5 | 192.0 |
|  |  | 3 | 7.0 | 3.5 | 11 | 31 | 113.2 | 202.7 | 183.0 |
|  |  | 4 | 6.6 | 3.5 | 16 | 41 | 154.0 | 109.5 | 181.0 |
|  |  | 5 | 6.5 | 3.0 | 16 | 31 | 110.6 | 149.3 | 168.0 |
|  |  | 6 | 6.7 | 2.5 | 16 | 31 | 94.1 | 125.6 | 166.2 |
|  |  | 7 | 6.4 | 3.0 | 11 | 31 | 145.4 | 220.9 | 198.4 |
|  |  | 8 | 6.7 | 2.5 | 16 | 31 | 104.9 | 172.9 | 211.2 |
|  |  | 9 | 6.4 | 3.0 | 16 | 31 | 111.0 | 261.5 | 192.3 |
|  |  | 10 | 6.1 | 2.5 | 16 | 31 | 135.7 | 203.3 | 221.8 |
|  |  | 11 | 7.0 | 2.5 | 16 | 21 | 57.0 | 89.5 | 182.9 |
|  |  | 12 | 6.8 | 3.0 | 16 | 31 | 93.3 | 117.1 | 151.5 |
|  |  | 13 | 6.8 | 3.5 | 11 | 41 | 162.4 | 184.7 | 228.6 |
|  |  | 14 | 6.3 | 2.5 | 16 | 21 | 102.0 | 193.4 | 218.1 |
|  |  | 15 | 6.7 | 2.5 | 16 | 21 | 118.9 | 164.9 | 171.5 |

**Fig. S2:**
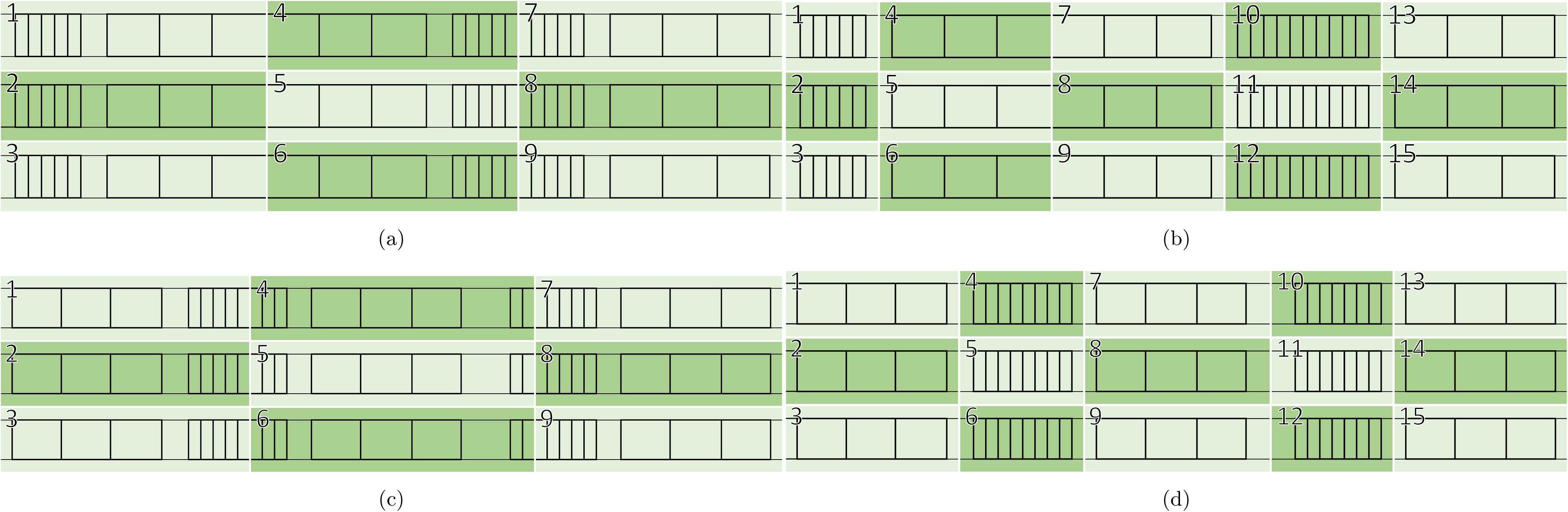
Location of soil sample collection in Langenthal BE in 2019 (a) and 2022 (b) and in Eschenbach LU in 2019 (c) and 2022 (d). For each of the numbered areas (i.e., 9 areas in 2019 and 15 areas in 2022 per location), one sample has been analysed which contained 10 subsamples collected along a diagonal of the area.

**Table S6:** Settings and quality of forage quality calibrations of dry matter (DM), acid detergent fibre (ADF), neutral detergent fibre (NDF), acid detergent lignin (ADL), crude protein (CP), non-fibre carbohydrates (NFC), ash, crude fat (CF), soluble protein (SP), and sulphur. The following pretreatments have been applied: first derivative using Savitzky-Golay (dg1), normalisation by closure (ncl), standard normal variate (SNV), first derivative using BCAP (db1), absorbance logarithm (log), first derivative using Savitzky-Golay and gap2-filtering (dg1g2), normalisation to unit length (nle). Additionally, the appropriate number of primary and secondary principal components (PCs) were set. Calibration quality was assessed using the standard error of calibration (SEC) and prediction (SEP), the coefficient of determination (R^2^) of the calibration and validation, the average difference between predicted and actual values (bias), and the residual predictive deviation (RPD).

| Property | DM | ADF | NDF | ADL | CP | NFC | Ash | CF | SP | Sulphur |
| --- | --- | --- | --- | --- | --- | --- | --- | --- | --- | --- |
| **Reference values** |  |  |  |  |  |  |  |  |  |  |
| Mean | 90.58 | 22.66 | 36.57 | 3.80 | 16.26 | 23.63 | 9.95 | 4.17 | 35.02 | 0.26 |
| Min | 87.60 | 14.80 | 22.87 | 0.73 | 7.24 | 0.27 | 5.29 | 1.91 | 22.82 | 0.11 |
| Max | 92.90 | 34.76 | 48.72 | 11.92 | 31.45 | 42.11 | 37.71 | 8.01 | 57.28 | 0.57 |
| SD | 0.83 | 3.34 | 5.27 | 1.92 | 4.76 | 7.51 | 3.77 | 1.06 | 5.13 | 0.09 |
| **Calibration settings** |  |  |  |  |  |  |  |  |  |  |
| Wave length range [cm<sup>-1</sup>] | 4200-8800 | 4200-9300 | 4200-9000 | 4000-9000 | 4200-9000 | 4000-9000 | 4100-4600, 4800-9000 | 4000-9000 | 4200-8500 | 4000-9500 |
| Calibration set [count] | 366 | 342 | 339 | 336 | 378 | 339 | 375 | 333 | 362 | 345 |
| Validation set [count] | 138 | 162 | 165 | 165 | 126 | 162 | 120 | 156 | 139 | 159 |
| Excluded measurements [count] | 0 | 0 | 0 | 3 | 0 | 3 | 9 | 15 | 3 | 0 |
| Pretreatment (first/second) | dg1/ncl | ncl/SNV | SNV/db1 | log/dg1g2 | -/- | SNV/- | SNV/dg1 | db1/nle | db1/- | ncl/db1 |
| Primary/Secondary PCs [count] | 11/11 | 11/11 | 9/9 | 10/10 | 11/11 | 9/9 | 11/11 | 10/10 | 11/11 | 9/9 |
| **Calibration quality** |  |  |  |  |  |  |  |  |  |  |
| SEC [%] | 0.45 | 6.84 | 3.86 | 30.77 | 3.44 | 6.98 | 5.83 | 8.15 | 6.28 | 16.00 |
| SEP [%] | 0.56 | 7.24 | 4.54 | 30.77 | 3.75 | 8.34 | 7.54 | 9.35 | 6.60 | 17.44 |
| $R^2$ calibration | 0.76 | 0.79 | 0.92 | 0.62 | 0.98 | 0.96 | 0.94 | 0.86 | 0.81 | 0.83 |
| $R^2$ validation | 0.64 | 0.74 | 0.90 | 0.52 | 0.98 | 0.92 | 0.88 | 0.77 | 0.79 | 0.72 |
| Bias [%] | -0.09 | 0.74 | 0.39 | -0.82 | -0.74 | 0.55 | 1.91 | -0.88 | -0.57 | 0.59 |
| RPD | 1.64 | 2.04 | 3.18 | 1.64 | 7.80 | 3.81 | 5.03 | 2.72 | 2.22 | 2.08 |

**Fig. S3:**
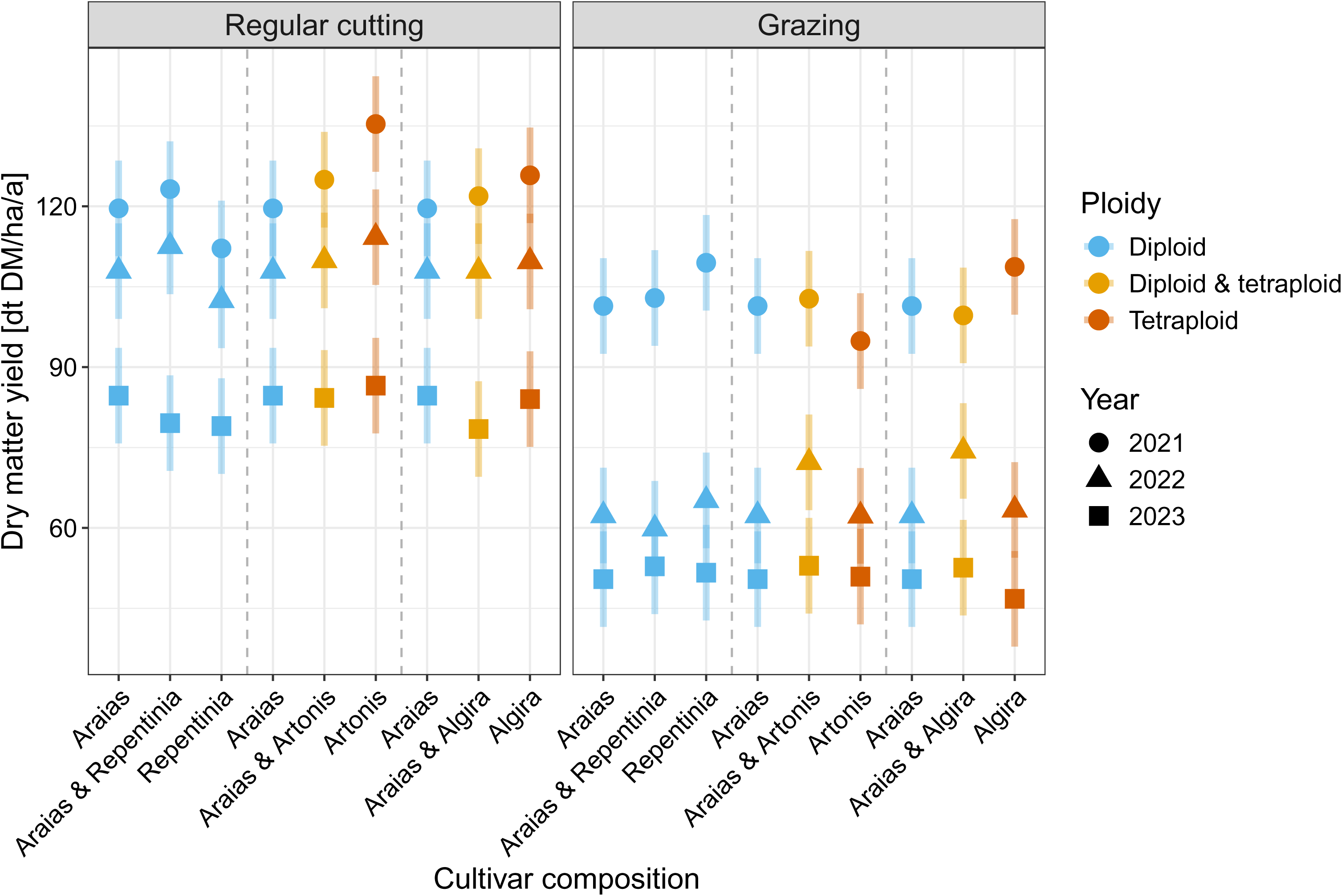
Estimated marginal means of yearly sum of dry matter yield per defoliation treatment, plot composition based on six *Lolium perenne* L. cultivars, and year, bars indicate 95% confidence intervals. Mixtures sown at a 50:50-ratio and the corresponding single-cultivar stands are shown in groups (separated by dashed lines) for direct comparison.

**Table S7:** Analysis of variance of the forage quality parameters, presenting the sum of squares (Sum Sq), the numerator and denominator degrees of freedom (df_Num_, df_Den_) of the *F* -test, the associated significance levels, and the partial R^2^ of each fixed effect, for crude protein (CP), non-fibre carbohydrates (NFC), neutral detergent fibre (NDF), acid detergent fibre (ADF), acid detergent lignin (ADL), and metabolisable energy (ME). In each row, the right-hand table omits the factor column and follows the same row order as the left-hand table.

| CP |  |  |  |  |  | NFC |  |  |  |  |  |
| --- | --- | --- | --- | --- | --- | --- | --- | --- | --- | --- | --- |
| Factor | Sum Sq | df<sub>Num</sub> | df<sub>Den</sub> | Partial $R^2$ | $F$-value | Sum Sq | df<sub>Num</sub> | df<sub>Den</sub> | Partial $R^2$ | $F$-value | |
| **Main effects** |  |  |  |  |  |  |  |  |  |  |  |
| Location | 10.80 | 1 | 4 | 0.073 | 1.15<sup>ns</sup> | 162.62 | 1 | 4 | 0.317 | 6.51<sup>ns</sup> |  |
| Plt. comp. | 293.52 | 8 | 105.5 | 0.020 | 3.92<sup>****</sup> | 2051.70 | 8 | 105.5 | 0.050 | 10.26<sup>****</sup> |  |
| Def. trtm. | 1336.29 | 2 | 10.5 | 0.903 | 71.35<sup>****</sup> | 1107.69 | 2 | 10.4 | 0.752 | 22.16<sup>****</sup> |  |
| Defoliation | 16085.11 | 3 | 1231.2 | 0.525 | 572.80<sup>****</sup> | 67028.69 | 3 | 1231.1 | 0.633 | 894.20<sup>****</sup> |  |
| Year | 3355.82 | 2 | 238.9 | 0.187 | 179.25<sup>****</sup> | 6402.55 | 2 | 238.9 | 0.141 | 128.12<sup>****</sup> |  |
| Residuals | 15454.16 | 1651 |  |  |  | 41252.52 | 1651 |  |  |  |  |
| **Interactions** |  |  |  |  |  |  |  |  |  |  |  |
| Plt. comp.×def. trtm. | 63.50 | 13 | 105.1 | 0.004 | 0.52<sup>ns</sup> | 461.79 | 13 | 105.0 | 0.012 | 1.42<sup>ns</sup> |  |
| Plt. comp.×year | 104.79 | 16 | 239.9 | 0.007 | 0.70<sup>ns</sup> | 188.85 | 16 | 239.9 | 0.005 | 0.47<sup>ns</sup> |  |
| Def. trtm.×year | 750.59 | 4 | 239.7 | 0.049 | 20.05<sup>****</sup> | 2065.47 | 4 | 239.7 | 0.050 | 20.67<sup>****</sup> |  |
| Def. trtm.×defoliation | 754.04 | 6 | 1231.6 | 0.049 | 13.43<sup>****</sup> | 7767.88 | 6 | 1231.5 | 0.166 | 51.81<sup>****</sup> |  |
| Plt. c.×def. trtm.×yr | 75.60 | 26 | 238.8 | 0.005 | 0.31<sup>ns</sup> | 203.61 | 26 | 238.8 | 0.005 | 0.31<sup>ns</sup> |  |
| NDF |  |  |  |  |  | ADF |  |  |  |  |  |
| Factor | Sum Sq | df<sub>Num</sub> | df<sub>Den</sub> | Partial $R^2$ | $F$-value | Sum Sq | df<sub>Num</sub> | df<sub>Den</sub> | Partial $R^2$ | $F$-value | |
| **Main effects** |  |  |  |  |  |  |  |  |  |  |  |
| Location | 120.11 | 1 | 4 | 0.359 | 7.92<sup>*</sup> | 1.66 | 1 | 4 | 0.013 | 0.19<sup>ns</sup> |  |
| Plt. comp. | 1679.06 | 8 | 105.5 | 0.066 | 13.84<sup>****</sup> | 1134.37 | 8 | 105.5 | 0.078 | 16.52<sup>****</sup> |  |
| Def. trtm. | 1558.48 | 2 | 10.5 | 0.874 | 51.36<sup>****</sup> | 355.73 | 2 | 10.2 | 0.740 | 20.72<sup>****</sup> |  |
| Defoliation | 11703.05 | 3 | 1231.3 | 0.331 | 257.23<sup>****</sup> | 8556.74 | 3 | 1231.0 | 0.391 | 332.37<sup>****</sup> |  |
| Year | 2547.84 | 2 | 238.8 | 0.097 | 84.00<sup>****</sup> | 417.81 | 2 | 238.9 | 0.030 | 24.34<sup>****</sup> |  |
| Residuals | 25038.05 | 1651 |  |  |  | 14168.12 | 1651 |  |  |  |  |
| **Interactions** |  |  |  |  |  |  |  |  |  |  |  |
| Plt. comp.×def. trtm. | 310.73 | 13 | 105.1 | 0.013 | 1.58<sup>ns</sup> | 82.26 | 13 | 105.0 | 0.006 | 0.74<sup>ns</sup> |  |
| Plt. comp.×year | 82.21 | 16 | 239.9 | 0.003 | 0.34<sup>ns</sup> | 43.87 | 16 | 239.9 | 0.003 | 0.32<sup>ns</sup> |  |
| Def. trtm.×year | 1429.05 | 4 | 239.7 | 0.057 | 23.56<sup>****</sup> | 1831.01 | 4 | 239.7 | 0.121 | 53.34<sup>****</sup> |  |
| Def. trtm.×defoliation | 4085.63 | 6 | 1231.7 | 0.148 | 44.90<sup>****</sup> | 1279.04 | 6 | 1231.4 | 0.087 | 24.84<sup>****</sup> |  |
| Plt. c.×def. trtm.×yr | 101.68 | 26 | 238.8 | 0.004 | 0.26<sup>ns</sup> | 45.05 | 26 | 238.8 | 0.003 | 0.20<sup>ns</sup> |  |
| ADL |  |  |  |  |  | ME |  |  |  |  |  |
| Factor | Sum Sq | df<sub>Num</sub> | df<sub>Den</sub> | Partial $R^2$ | $F$-value | Sum Sq | df<sub>Num</sub> | df<sub>Den</sub> | Partial $R^2$ | $F$-value | |
| **Main effects** |  |  |  |  |  |  |  |  |  |  |  |
| Location | 2.51 | 1 | 4 | 0.151 | 2.52<sup>ns</sup> | 0.24 | 1 | 4 | 0.123 | 2.00<sup>ns</sup> |  |
| Plt. comp. | 109.53 | 8 | 105.7 | 0.067 | 13.74<sup>****</sup> | 20.87 | 8 | 105.4 | 0.101 | 21.70<sup>****</sup> |  |
| Def. trtm. | 11.34 | 2 | 10.2 | 0.441 | 5.69<sup>*</sup> | 12.35 | 2 | 10.2 | 0.877 | 51.35<sup>****</sup> |  |
| Defoliation | 3268.31 | 3 | 1212.2 | 0.682 | 1093.38<sup>****</sup> | 325.41 | 3 | 1228.4 | 0.636 | 902.30<sup>****</sup> |  |
| Year | 484.55 | 2 | 238.6 | 0.241 | 243.15<sup>****</sup> | 8.20 | 2 | 238.3 | 0.042 | 34.12<sup>****</sup> |  |
| Residuals | 1617.14 | 1623 |  |  |  | 198.00 | 1647 |  |  |  |  |
| **Interactions** |  |  |  |  |  |  |  |  |  |  |  |
| Plt. comp.×def. trtm. | 8.12 | 13 | 104.7 | 0.005 | 0.63<sup>ns</sup> | 2.04 | 13 | 105.0 | 0.011 | 1.31<sup>ns</sup> |  |
| Plt. comp.×year | 7.16 | 16 | 240.2 | 0.005 | 0.45<sup>ns</sup> | 1.21 | 16 | 239.7 | 0.006 | 0.63<sup>ns</sup> |  |
| Def. trtm.×year | 79.44 | 4 | 238.3 | 0.050 | 19.93<sup>****</sup> | 23.92 | 4 | 239.6 | 0.114 | 49.73<sup>****</sup> |  |
| Def. trtm.×defoliation | 257.05 | 6 | 1212.7 | 0.144 | 43.00<sup>****</sup> | 99.75 | 6 | 1228.8 | 0.348 | 138.30<sup>****</sup> |  |
| Plt. c.×def. trtm.×yr | 11.50 | 26 | 237.9 | 0.008 | 0.44<sup>ns</sup> | 0.54 | 26 | 238.7 | 0.003 | 0.17<sup>ns</sup> |  |

ns: $P > 0.05$ , \*: $P \leq 0.05$ , \*\*: $P \leq 0.01$ , \*\*\*: $P \leq 0.001$ , \*\*\*\*: $P \leq 0.0001$ . Partial $R^2$ values are only directly comparable within the same denominator range (e.g. location and defoliation treatment on ~4–10 df; cultivar-related terms on ~105 df; year-related terms on ~239–265 df).

**Table S8:** Percentage of total random-effect variance attributable to each component, for the yearly sum of dry matter yield (DMY), neutral detergent fibre (NDF), acid detergent fibre (ADF), acid detergent lignin (ADL), crude protein (CP), non-fibre carbohydrates (NFC), and metabolisable energy (ME).

| Component | DMY | NDF | ADF | ADL | CP | NFC | ME |
| --- | --- | --- | --- | --- | --- | --- | --- |
| Plot-year [%] | – | 0.0 | 0.0 | 0.0 | 0.0 | 0.0 | 0.0 |
| Plot [%] | 6.4 | 0.0 | 0.0 | 0.0 | 0.0 | 0.0 | 0.0 |
| Whole plot (def. trtm.) [%] | 9.0 | 2.3 | 5.5 | 8.1 | 2.5 | 3.5 | 8.6 |
| Block (within location) [%] | 0.2 | 0.0 | 0.0 | 0.0 | 0.0 | 0.0 | 0.0 |
| Residual [%] | 84.5 | 97.7 | 94.5 | 91.9 | 97.5 | 96.5 | 91.4 |

–: not applicable; the plot-year random effect accounts for the four repeated defoliation measurements within a plot-year and is therefore only present in the forage quality models, which are based on individual defoliations rather than the yearly sum.

**Table S9:** Diploid minus tetraploid difference in dry matter yield (DMY) and each forage quality parameter, within each defoliation treatment (C: regular cutting, G: grazing, SG: simulated grazing). Positive values indicate diploid cultivars higher; negative values indicate tetraploid cultivars higher.

| Parameter | Defoliation treatment | Estimate | SE | 95% CI | Sig. |
| --- | --- | --- | --- | --- | --- |
| DMY [dt DM/ha/a] | C | -7.01 | 2.17 | [-11.32, -2.70] | ** |
|  | G | 3.72 | 2.17 | [-0.59, 8.03] | ns |
|  | SG | -1.67 | 2.17 | [-5.98, 2.63] | ns |
| NDF [%] | C | 2.88 | 0.38 | [2.13, 3.64] | **** |
|  | G | 1.89 | 0.39 | [1.12, 2.65] | **** |
|  | SG | 1.44 | 0.38 | [0.69, 2.18] | *** |
| ADF [%] | C | 2.12 | 0.29 | [1.55, 2.68] | **** |
|  | G | 1.29 | 0.29 | [0.72, 1.86] | **** |
|  | SG | 1.64 | 0.28 | [1.08, 2.20] | **** |
| ADL [%] | C | 0.56 | 0.10 | [0.36, 0.75] | **** |
|  | G | 0.44 | 0.10 | [0.24, 0.64] | **** |
|  | SG | 0.56 | 0.10 | [0.37, 0.75] | **** |
| CP [%] | C | -0.11 | 0.30 | [-0.70, 0.49] | ns |
|  | G | -0.55 | 0.30 | [-1.15, 0.05] | ns |
|  | SG | 0.08 | 0.30 | [-0.51, 0.66] | ns |
| NFC [%] | C | -2.68 | 0.49 | [-3.65, -1.71] | **** |
|  | G | -1.00 | 0.49 | [-1.98, -0.02] | * |
|  | SG | -1.70 | 0.48 | [-2.66, -0.74] | *** |
| ME [MJ/kg] | C | -0.31 | 0.03 | [-0.38, -0.24] | **** |
|  | G | -0.18 | 0.03 | [-0.25, -0.11] | **** |
|  | SG | -0.22 | 0.03 | [-0.28, -0.15] | **** |

ns: $P > 0.05$ , \*: $P \leq 0.05$ , \*\*: $P \leq 0.01$ , \*\*\*: $P \leq 0.001$ , \*\*\*\*: $P \leq 0.0001$ .

**Table S10:** Difference between defoliation treatments in the diploid-tetraploid contrast (i.e., the ploidy × defoliation treatment interaction), for dry matter yield (DMY) and each forage quality parameter (C: regular cutting, G: grazing, SG: simulated grazing). A positive value indicates a larger diploid-minus-tetraploid difference under the first-named defoliation treatment.

| Parameter | C vs G | C vs SG | G vs SG |
| --- | --- | --- | --- |
| DMY [dt DM/ha/a] | -10.73<sup>***</sup> | -5.34<sup>ns</sup> | 5.39<sup>ns</sup> |
| NDF [%] | 1.04<sup>ns</sup> | 1.47<sup>**</sup> | 0.44<sup>ns</sup> |
| ADF [%] | 0.82<sup>*</sup> | 0.47<sup>ns</sup> | -0.35<sup>ns</sup> |
| ADL [%] | 0.12<sup>ns</sup> | 0.00<sup>ns</sup> | -0.11<sup>ns</sup> |
| CP [%] | 0.41<sup>ns</sup> | -0.19<sup>ns</sup> | -0.60<sup>ns</sup> |
| NFC [%] | -1.65<sup>*</sup> | -0.98<sup>ns</sup> | 0.66<sup>ns</sup> |
| ME [MJ/kg] | -0.13<sup>**</sup> | -0.09<sup>ns</sup> | 0.04<sup>ns</sup> |

ns: $P > 0.05$ , \*: $P \leq 0.05$ , \*\*: $P \leq 0.01$ , \*\*\*: $P \leq 0.001$ .

*Reading examples:* For DMY, C vs G = -10.73 means the diploid-minus-tetraploid difference in DMY was 10.73 dt DM/ha/a more negative under regular cutting than under grazing (Supplementary Table S9: -7.01 under C, +3.72 under G); that is, the tetraploid yield advantage over diploid cultivars was much larger under regular cutting, whereas diploid cultivars performed slightly better under grazing. For NDF, where diploid cultivars had higher values than tetraploid cultivars under all three defoliation treatments (Supplementary Table S9), the positive C vs SG = 1.47 means this diploid-tetraploid difference was larger under regular cutting than under simulated grazing.

**Table S11:**
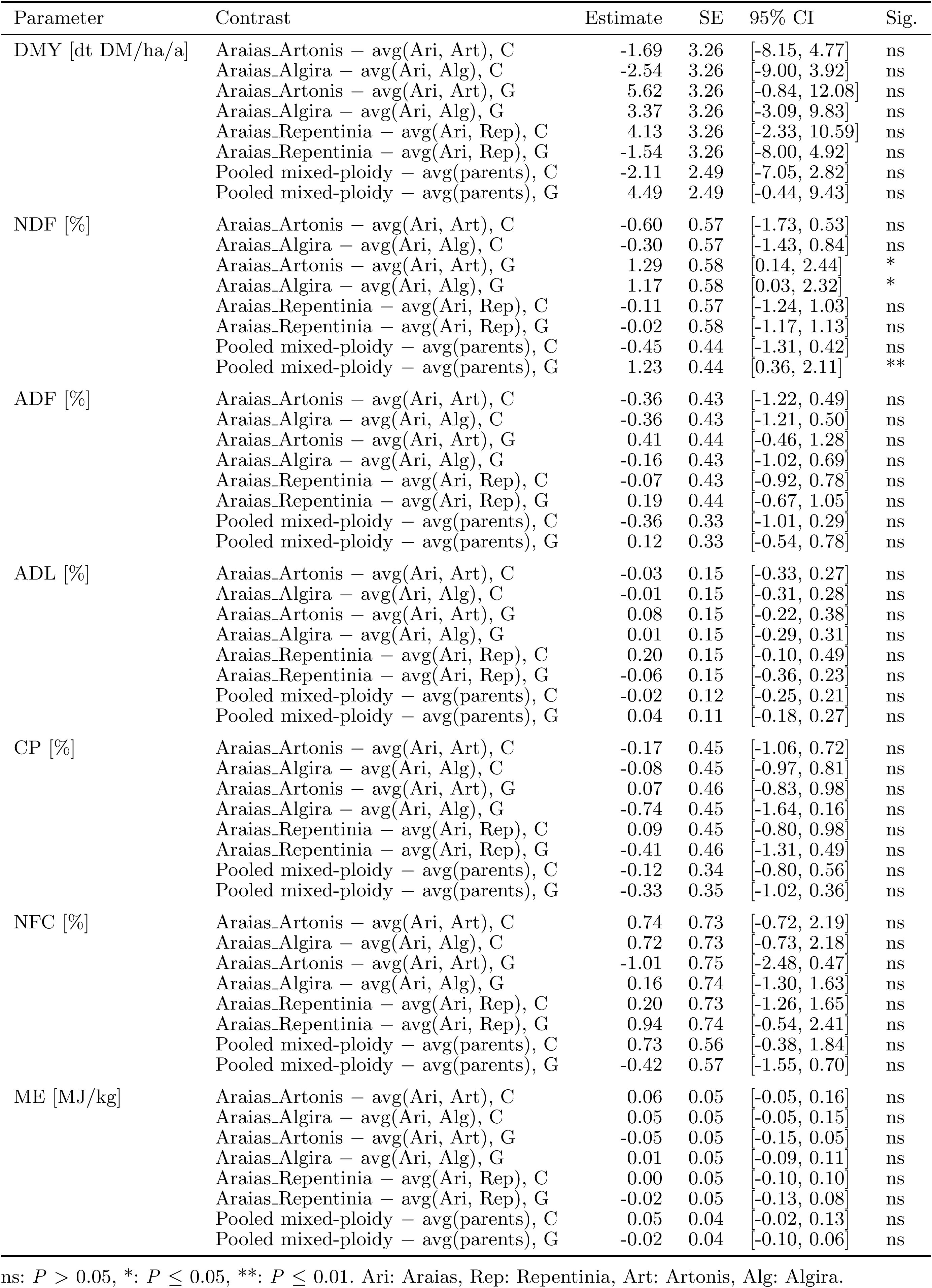
Deviation of each mixture from the average of its component cultivars, for dry matter yield (DMY) and each forage quality parameter, under regular cutting (C) and grazing (G).

| Parameter | Contrast | Estimate | SE | 95% CI | Sig. |
| --- | --- | --- | --- | --- | --- |
| DMY [dt DM/ha/a] | Araias_Artonis – avg(Ari, Art), C | -1.69 | 3.26 | [-8.15, 4.77] | ns |
|  | Araias_Algira – avg(Ari, Alg), C | -2.54 | 3.26 | [-9.00, 3.92] | ns |
|  | Araias_Artonis – avg(Ari, Art), G | 5.62 | 3.26 | [-0.84, 12.08] | ns |
|  | Araias_Algira – avg(Ari, Alg), G | 3.37 | 3.26 | [-3.09, 9.83] | ns |
|  | Araias_Repentinia – avg(Ari, Rep), C | 4.13 | 3.26 | [-2.33, 10.59] | ns |
|  | Araias_Repentinia – avg(Ari, Rep), G | -1.54 | 3.26 | [-8.00, 4.92] | ns |
|  | Pooled mixed-ploidy – avg(parents), C | -2.11 | 2.49 | [-7.05, 2.82] | ns |
|  | Pooled mixed-ploidy – avg(parents), G | 4.49 | 2.49 | [-0.44, 9.43] | ns |
| NDF [%] | Araias_Artonis – avg(Ari, Art), C | -0.60 | 0.57 | [-1.73, 0.53] | ns |
|  | Araias_Algira – avg(Ari, Alg), C | -0.30 | 0.57 | [-1.43, 0.84] | ns |
|  | Araias_Artonis – avg(Ari, Art), G | 1.29 | 0.58 | [0.14, 2.44] | * |
|  | Araias_Algira – avg(Ari, Alg), G | 1.17 | 0.58 | [0.03, 2.32] | * |
|  | Araias_Repentinia – avg(Ari, Rep), C | -0.11 | 0.57 | [-1.24, 1.03] | ns |
|  | Araias_Repentinia – avg(Ari, Rep), G | -0.02 | 0.58 | [-1.17, 1.13] | ns |
|  | Pooled mixed-ploidy – avg(parents), C | -0.45 | 0.44 | [-1.31, 0.42] | ns |
|  | Pooled mixed-ploidy – avg(parents), G | 1.23 | 0.44 | [0.36, 2.11] | ** |
| ADF [%] | Araias_Artonis – avg(Ari, Art), C | -0.36 | 0.43 | [-1.22, 0.49] | ns |
|  | Araias_Algira – avg(Ari, Alg), C | -0.36 | 0.43 | [-1.21, 0.50] | ns |
|  | Araias_Artonis – avg(Ari, Art), G | 0.41 | 0.44 | [-0.46, 1.28] | ns |
|  | Araias_Algira – avg(Ari, Alg), G | -0.16 | 0.43 | [-1.02, 0.69] | ns |
|  | Araias_Repentinia – avg(Ari, Rep), C | -0.07 | 0.43 | [-0.92, 0.78] | ns |
|  | Araias_Repentinia – avg(Ari, Rep), G | 0.19 | 0.44 | [-0.67, 1.05] | ns |
|  | Pooled mixed-ploidy – avg(parents), C | -0.36 | 0.33 | [-1.01, 0.29] | ns |
|  | Pooled mixed-ploidy – avg(parents), G | 0.12 | 0.33 | [-0.54, 0.78] | ns |
| ADL [%] | Araias_Artonis – avg(Ari, Art), C | -0.03 | 0.15 | [-0.33, 0.27] | ns |
|  | Araias_Algira – avg(Ari, Alg), C | -0.01 | 0.15 | [-0.31, 0.28] | ns |
|  | Araias_Artonis – avg(Ari, Art), G | 0.08 | 0.15 | [-0.22, 0.38] | ns |
|  | Araias_Algira – avg(Ari, Alg), G | 0.01 | 0.15 | [-0.29, 0.31] | ns |
|  | Araias_Repentinia – avg(Ari, Rep), C | 0.20 | 0.15 | [-0.10, 0.49] | ns |
|  | Araias_Repentinia – avg(Ari, Rep), G | -0.06 | 0.15 | [-0.36, 0.23] | ns |
|  | Pooled mixed-ploidy – avg(parents), C | -0.02 | 0.12 | [-0.25, 0.21] | ns |
|  | Pooled mixed-ploidy – avg(parents), G | 0.04 | 0.11 | [-0.18, 0.27] | ns |
| CP [%] | Araias_Artonis – avg(Ari, Art), C | -0.17 | 0.45 | [-1.06, 0.72] | ns |
|  | Araias_Algira – avg(Ari, Alg), C | -0.08 | 0.45 | [-0.97, 0.81] | ns |
|  | Araias_Artonis – avg(Ari, Art), G | 0.07 | 0.46 | [-0.83, 0.98] | ns |
|  | Araias_Algira – avg(Ari, Alg), G | -0.74 | 0.45 | [-1.64, 0.16] | ns |
|  | Araias_Repentinia – avg(Ari, Rep), C | 0.09 | 0.45 | [-0.80, 0.98] | ns |
|  | Araias_Repentinia – avg(Ari, Rep), G | -0.41 | 0.46 | [-1.31, 0.49] | ns |
|  | Pooled mixed-ploidy – avg(parents), C | -0.12 | 0.34 | [-0.80, 0.56] | ns |
|  | Pooled mixed-ploidy – avg(parents), G | -0.33 | 0.35 | [-1.02, 0.36] | ns |
| NFC [%] | Araias_Artonis – avg(Ari, Art), C | 0.74 | 0.73 | [-0.72, 2.19] | ns |
|  | Araias_Algira – avg(Ari, Alg), C | 0.72 | 0.73 | [-0.73, 2.18] | ns |
|  | Araias_Artonis – avg(Ari, Art), G | -1.01 | 0.75 | [-2.48, 0.47] | ns |
|  | Araias_Algira – avg(Ari, Alg), G | 0.16 | 0.74 | [-1.30, 1.63] | ns |
|  | Araias_Repentinia – avg(Ari, Rep), C | 0.20 | 0.73 | [-1.26, 1.65] | ns |
|  | Araias_Repentinia – avg(Ari, Rep), G | 0.94 | 0.74 | [-0.54, 2.41] | ns |
|  | Pooled mixed-ploidy – avg(parents), C | 0.73 | 0.56 | [-0.38, 1.84] | ns |
|  | Pooled mixed-ploidy – avg(parents), G | -0.42 | 0.57 | [-1.55, 0.70] | ns |
| ME [MJ/kg] | Araias_Artonis – avg(Ari, Art), C | 0.06 | 0.05 | [-0.05, 0.16] | ns |
|  | Araias_Algira – avg(Ari, Alg), C | 0.05 | 0.05 | [-0.05, 0.15] | ns |
|  | Araias_Artonis – avg(Ari, Art), G | -0.05 | 0.05 | [-0.15, 0.05] | ns |
|  | Araias_Algira – avg(Ari, Alg), G | 0.01 | 0.05 | [-0.09, 0.11] | ns |
|  | Araias_Repentinia – avg(Ari, Rep), C | 0.00 | 0.05 | [-0.10, 0.10] | ns |
|  | Araias_Repentinia – avg(Ari, Rep), G | -0.02 | 0.05 | [-0.13, 0.08] | ns |
|  | Pooled mixed-ploidy – avg(parents), C | 0.05 | 0.04 | [-0.02, 0.13] | ns |
|  | Pooled mixed-ploidy – avg(parents), G | -0.02 | 0.04 | [-0.10, 0.06] | ns |

ns: $P > 0.05$ , \*: $P \leq 0.05$ , \*\*: $P \leq 0.01$ . Ari: Araias, Rep: Repentinia, Art: Artonis, Alg: Algira.

